# Desmosomal Cadherin Tension Loss in Pemphigus Vulgaris Mediated by the Inhibition of Active RhoA at Cell-Cell Adhesions

**DOI:** 10.1101/2024.05.03.592394

**Authors:** Xiaowei Jin, Jordan Rosenbohm, Amir Ostadi Moghaddam, Eunju Kim, Kristina Seiffert-Sinha, Merced Leiker, Haiwei Zhai, Sindora R. Baddam, Grayson Minnick, Yucheng Huo, Bahareh Tajvidi Safa, James K. Wahl, Fanben Meng, Changjin Huang, Jung Yul Lim, Daniel E. Conway, Animesh A. Sinha, Ruiguo Yang

## Abstract

Binding of autoantibodies to keratinocyte surface antigens, primarily desmoglein 3 (Dsg3) of the desmosomal complex, leads to the dissociation of cell-cell adhesion in the blistering disorder pemphigus vulgaris (PV). After the initial disassembly of desmosomes, cell-cell adhesions actively remodel in association with the cytoskeleton and focal adhesions. Growing evidence highlights the role of adhesion mechanics and mechanotransduction at cell-cell adhesions in this remodeling process, as their active participation may direct autoimmune pathogenicity. However, a large part of the biophysical transformations after antibody binding remains underexplored. Specifically, it is unclear how tension in desmosomes and cell-cell adhesions changes in response to antibodies, and how the altered tensional states translate to cellular responses. Here, we showed a tension loss at Dsg3 using fluorescence resonance energy transfer (FRET)-based tension sensors, a tension loss at the entire cell-cell adhesion, and a potentially compensatory increase in junctional traction force at cell-extracellular matrix adhesions after PV antibody binding. Further, our data indicate that this tension loss is mediated by the inhibition of RhoA at cell-cell contacts, and the extent of RhoA inhibition may be crucial in determining the severity of pathogenicity among different PV antibodies. More importantly, this tension loss can be partially restored by altering actomyosin based cell contractility. Collectively, these findings provide previously unattainable details in our understanding of the mechanisms that govern cell-cell interactions under physiological and autoimmune conditions, which may open the window to entirely new therapeutics aimed at restoring physiological balance to tension dynamics that regulates the maintenance of cell-cell adhesion.

## INTRODUCTION

Desmosomes are cell-cell adhesions that play a critical role in maintaining the structural integrity of tissues. They are the main target in pemphigus vulgaris (PV), in which autoantibodies (autoAbs) recognize desmosomal proteins, desmoglein 3 (Dsg3) and in some cases Dsg1 as well, leading to the breakdown of cell-cell adhesion and the formation of blisters. The exact mechanisms underlying desmosome dissociation in PV are poorly understood, but current research suggests that autoAbs trigger a signaling cascade that ultimately results in desmosome disassembly.^1^ The suppression of autoAb generation or their removal from circulation is currently the primary approach to therapy in PV.^2^ However, immunosuppression comes with many detrimental side effects, and a patient’s comorbidities can restrict treatment options.^3^ A deeper understanding of the biophysical signaling pathways induced by PV autoAb binding holds the potential to reveal new therapeutic targets. These pathways involve molecules that govern the overall cell contractility and tension at the cell-cell junctions, including E-cadherin,^4^ Rho GTPase,^5^ and Myosin II.^6^ Recent literature suggests that the desmosome disassembly process that ultimately leads to blister formation significantly modifies cell mechanics.^7^ In response, cell-cell adhesions actively remodel in association with the cytoskeleton and other adhesion structures to regulate their biophysical behavior.^8^ Growing evidence now highlights the role of adhesion mechanics and mechanotransduction in this adaptation process.^9^ However, the impact of changes in junction tension on altering the cause of adhesion dissociation is unclear.

Adherens junctions, the E-cadherin based adhesion complex, are closely linked to desmosomes and can also play a role in desmosome dissociation in PV. It has been shown that tissue expression of E-cadherin in PV patients varies between phases of disease activity and remission, and that PV patients can carry low levels of anti-E-cadherin antibodies (Abs).^10,11^ Furthermore, overexpression of Dsg3 results in not only a reduced level of E-cadherin but its colocalization with the E-cadherin-catenin complex of adherens junctions, suggesting that Dsg3 helps regulate adherens junction formation.^12^ In fact, it has been proposed that E-cadherin may play a compensatory role in maintaining cell-cell adhesion in the absence of functional desmosomes.^13^ However, the overall change in junction stress through this potential mechanical compensation after PV Ab binding has not been delineated. In addition, Rho GTPases, specifically RhoA, have been implicated in the downstream signaling pathways activated by anti-Dsg Abs, leading to alterations in actin cytoskeleton organization and increased activity of cell contractility.^5^ For instance, inhibition of RhoA activity in the epidermis in an inducible mouse model led to the formation of skin blisters, suggesting that RhoA activity is necessary for maintaining the integrity of desmosomal adhesion in the skin.^5^ It is yet to be seen how RhoA activities are regulated in association with junction tension and the overall cell contractility.

We have previously investigated the role of mechanical stress in modulating the dissociation of cell-cell adhesion induced by anti-Dsg3 Abs in PV.^14^ We showed that increasing mechanical stress on cells through stretching or shear forces could reduce the disruptive effects of anti-Dsg3 Abs on cell adhesion, suggesting that external mechanical stress at the junction may have a protective effect on cell-cell adhesions. In the current study, we report the use of fluorescence resonance energy transfer (FRET) sensors to further characterize biophysical transformations after PV Ab binding. These sensors allow for visualization of changes in molecular-scale distances, enabling the study of the dynamics of the proteins they are incorporated in. We show reduced tension within desmosomal cadherins and cell-cell adhesions as a whole after PV Ab binding, suggesting the dissociation at desmosomal cadherins is accompanied by tension loss. Concurrently, we observe increasing junctional traction stress at cell-extracellular matrix (ECM) adhesions from traction force microscopy, indicating potential mechanical compensation between cell-ECM and cell-cell adhesions. Importantly, inactivation of RhoA in cells treated with PV autoAbs is observed using a RhoA FRET sensor, suggesting that the mechanotransduction pathway involves RhoA signaling. Finally, we find that direct stimulation of RhoA reverses many detrimental effects of PV autoAb binding and restores tension across cell-cell and cell-ECM adhesions, highlighting its central role in the regulation of PV pathogenesis.

## RESULTS

### Desmosomes are destabilized by PV-associated Abs

We first characterized the dissociation of PV-associated Abs on HaCaT cells, a keratinocyte cell line. HaCaT cells were grown to 80% confluency with low calcium media in petri dishes and were cultured in high calcium media overnight before antibody treatments. HaCaT cells were treated with two types of PV-associated Abs: AK23, a commercial mouse-derived anti-human Dsg3 monoclonal Ab (mAb), and PVIgG, polyclonal Abs purified from anti Dsg3-containing patient serum. AK23 at 10 μg/mL concentrations, and PVIgG at 2 μg/mL, lead to a dispersed distribution pattern of Dsg3 (white arrow) compared with the tightly packed Dsg3 along the cell-cell contacts in controls at both 4h and 24h timepoints (Figure 1A). This scattered pattern, observed in both AK23 and PVIgG treated samples, is quantified by the mean width of Dsg3, using the value of the full width at half maximum (FWHM) of the immunofluorescence intensity profile along a line drawn between two neighboring cells (Figure S1).^15^ The FWHM, as described in a previous report,^15^ can be used to compare the localization of molecules and indicate how tightly they are associated with the junction. Compared to controls, the FWHM did not change for 2 μg/mL of AK23, while it increased around 2.5-fold for 10 μg/mL in AK23 groups at 4h and this increment lasted at 24h. For PVIgG, compared to controls, width increased around 4-fold and 5-fold at 4h and 24h, respectively (Figure 1B). The time-series data indicates that PV Abs induced sustained desmosome dissociation with a dosage-dependent effect, and that PVIgG is more potent and led to a more significant dissociation compared with AK23. We further confirmed that potential cross reaction between mouse-derived monoclonal antibodies and AK23 did not influence the reported values of FWHM (Figure S2). It is worth mentioning that we performed titration experiments which showed that dosages of 1 μg/mL and 2 μg/mL for AK23 resulted in loss of intercellular adhesion using keratinocyte dissociation assays, while PVIgG showed adhesion loss at 100 μg/mL (Figure S3 A and C).

**Figure 1.**
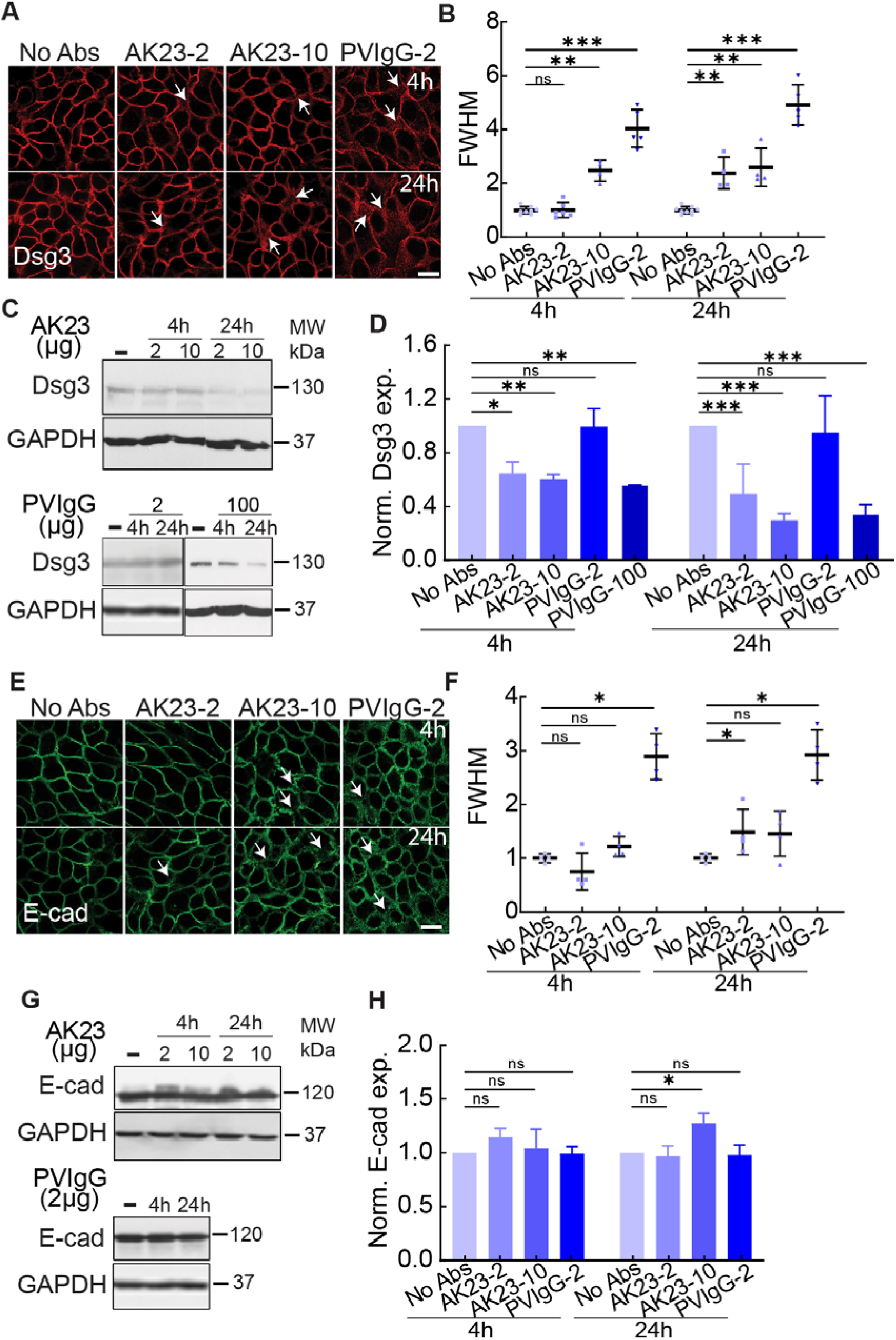
PV Abs induced junctional dissociation. Junctional cohesion was quantified with immunostaining images and FWHM, which represents the mean width of cell–cell junctions, and Dsg3 and E-cadherin expressions were evaluated by western blot in HaCaT cells exposed to different dosages of AK23 and PVIgG at multiple time points. (A) Dsg3 immunofluorescent images in groups of no treatment (No Abs), 2 µg/mL AK23 (AK23-2), 10 µg/mL AK23 (AK23-10), and 2 µg/mL PVIgG (PVIgG-2) at 4h and 24h. (B) Quantitative FWHM of Dsg3 with PV Ab treatments at 4h and 24h. All values are mean ± SD (n≥30). (C, D) Western blot results of Dsg3 with same treatments plus 100 µg/mL PVIgG (PVIgG-100) at 4h and 24h. All values are mean ± SD (n≥3). (E) E-cadherin immunofluorescent images in groups of no treatment (No Abs), 2 µg/mL AK23 (AK23-2), 10 µg/mL AK23 (AK23-10), and 2 µg/mL PVIgG (PVIgG-2) at 4h and 24h. (F) Quantitative FWHM of E-cadherin with PV Ab treatments at 4h and 24h. All values are mean ± SD (n≥30). (G, H) Western blot results of Dsg3 with same treatments at 4h and 24h. All values are mean ± SD (n≥3). * p<0.05, ** p<0.01, *** p<0.005, ns, p>0.05. Scale bar: 10 µm. Three repeats for all conditions.

The observed junctional morphology and increasing Dsg3 distribution width induced by PV-associated Ab treatment was accompanied by a downregulation of Dsg3 expression (Figure 1C and D). Both AK23 concentrations suppressed Dsg3 levels at 4h and 24h. Interestingly, even though PVIgG at 2 μg/mL causes desmosome destabilization observed from Dsg3 imaging, it did not lead to any significant change in overall Dsg3 expression, while a higher concentration of PVIgG (100 μg/mL) reduced Dsg3 expression significantly for both 4h and 24h timepoints, suggesting that different destabilization mechanisms may be at work for the two PV Abs.^16^ This difference could be the result of the two PV Abs interacting with different extracellular domains of Dsg3 or the added effects of PVIgG interfering with Dsg1^1,17^ and/or other potential targets, described in the literature before.^18,19^ The difference was also supported by a smaller reduction in anisotropy of intermediate filaments (IF) in PVIgG (2 μg/mL) compared to AK23 (2 μg/mL), indicating PVIgG induces less reorientation of IFs compared to AK23 (Figure S4). In addition, the difference between the expression of Dsg3 for AK23 treatments of 2 μg/mL and 10 μg/mL was larger at 24h compared to 4h, which suggested that endocytosis of Dsg3 in the cytoplasm is both time and dosage dependent.^20^

We next examined whether PV Abs result in destabilization of adherens junctions, which are a critical anchoring point for the actin cytoskeleton and contribute to overall cell contractility. First, the distribution of E-cadherin at the cell-cell contact was disrupted with PVIgG as shown by a significant increase in FWHM (Figure 1E and F), indicating PVIgG also destabilized the adherens junctions. Interestingly, we observed a decreased FWHM for 2 μg/mL in AK23 groups at 4h. Furthermore, E-cadherin protein expression exhibited no statistically significant changes for both AK23 concentrations at 4h (Figure 1G and H), but increased by 40% after 24h of AK23 at the higher concentration of 10 µg/ml, indicating the synergy between desmosomes and adherens junctions. We also observed an influence of PV Abs on the actin cytoskeleton, which was less tightly associated with the junction for AK23 at 24h and PVIgG at both timepoints (Figure S5). The synergy between these two adhesion types was further supported by the fact that significant changes to adherens junction-associated molecules only occur under conditions that induced significant changes to desmosome-associated molecules, indicating changes to adherens junction may follow changes to desmosomes (Figure 1, Figure S4 and S5).

### PV-associated Abs reduce Dsg3 tension at cell-cell adhesions as measured by FRET tension sensors

After confirming that PV Abs can destabilize desmosome structure, we next tested whether this disruption was accompanied by changes in desmosome tension. We theorized that tension across Dsg3 would change in response to changes in intercellular forces mediated by cell-cell junctions and intracellular forces mediated by cytoskeleton contractility. To quantify this tension, we used a FRET-based Dsg3 tension sensor. The Dsg3 sensor was created by inserting a tension sensor module (TSMod) with two fluorophores, an mTFP donor and a Venus acceptor (Figure 2A, Dsg3 TS). The energy transfer between the donor/acceptor pair is measured as a FRET ratio, calculated by dividing the fluorescent intensity of the acceptor by that of the donor (Figure S6). Since the FRET ratio is sensitive to distance between the fluorophore pair, a decreasing FRET ratio correlates with an increasing distance, suggesting higher tension; conversely, an increasing ratio is an indicator for reduced tension. A tailless sensor with the depletion of the cytoplasmic tail was used for control (Figure 2A, Dsg3 TL). When transfected into HaCaT cells, both sensors localized to cell-cell adhesions, as evidenced by their colocalization with desmoplakin (DP) (Figure 2B). Furthermore, immunostaining images of Dsg3 confirmed the recombination of the FRET sensor and Dsg3 cDNA, while immunogold electron microscopy images of Venus showed the localization of the Dsg3 sensor within the cell-cell contact (Figure 2B and C). Finally, a Triton X-100 fractionation assay was used to further demonstrate that the sensor integrates into desmosomes at the cell-cell contact similarly to endogenous Dsg3 (Figure S7).

**Figure 2.**
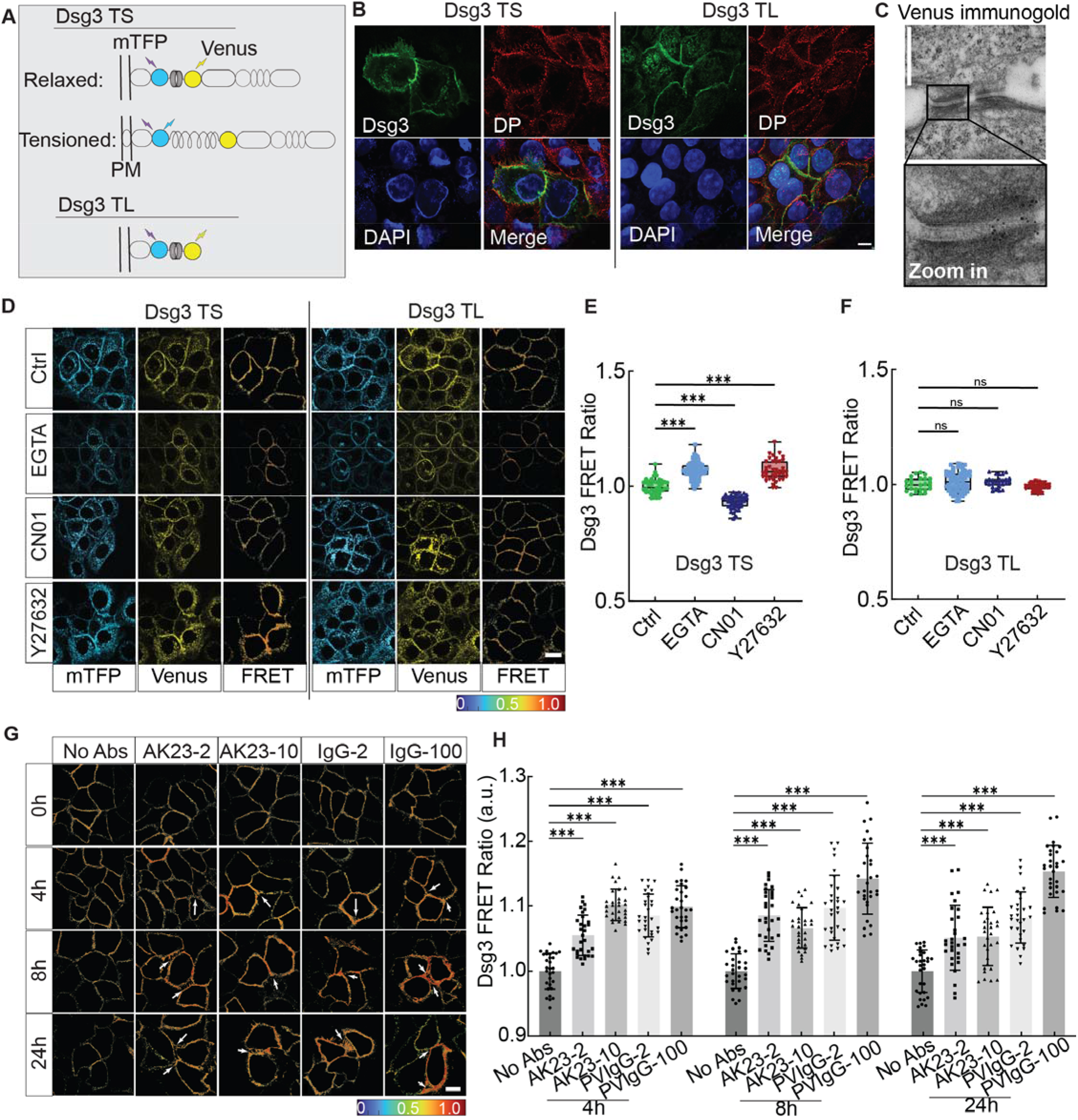
Dsg3 in response to inter- and intra-cellular force changes. (A) Schematic diagram of Dsg3 tension sensor (TS) and tailless control sensor (TL). TSMod was inserted into the intracellular domain of Dsg3. (B) Immunostaining images of desmoplakin (DP) for HaCaT cells expressing Dsg3 TS or TL. Cells were immunostained for DP to check for the co-expression with sensors and nuclei were labeled with DAPI. Scale bar: 10 µm. (C) Immunogold image for HaCaT cells expressing the Dsg3 tension sensor. Positive signals were observed at desmosomes, suggesting that Dsg3 tension sensors incorporated into desmosomal cadherins. Scale bar: 500 nm. (D) Fluorescent images of donor (mTFP) and acceptor (Venus) and heatmap images of FRET ratio when Dsg3 TS/Dsg3 TL expressed HaCaT cells were exposed to EGTA for 30 min, 5 mM RhoA activator (CN01) for 30 min, ROCK inhibitor (Y27632) for 12h. Scale bar: 10 µm. (E, F) Quantitative analysis of FRET ratio shown in D. (G) Heatmap images of FRET ratio in HaCaT cells treated with the following conditions: untreated control (No Abs), 2 µg/mL AK23 (AK23-2), 10 µg/mL AK23 (AK23-10), 2 µg/mL PVIgG (PVIgG-2) and 100 µg/mL PVIgG (PVIgG-100) at 0h, 4h, 8h, and 24h. Scale bar: 10 µm. (H) Quantitative analysis of corresponding FRET ratio. All values are mean ± SD (n≥30). * p<0.05, ** p<0.01, *** p<0.005, ns, p>0.05. Three repeats for all conditions.

To test whether Dsg3 responded to the intercellular tension loss, EGTA, a calcium chelator which induces the dissociation of cell junctions by interfering with the binding at the EC1, was used at a concentration of 5 mM for 30 min. We observed an increase in the FRET ratio for Dsg3 FRET tension sensor while no significant FRET ratio changes were detected for Dsg3 TL (Figure 2D-F). Moreover, to evaluate if the sensor also responded to intracellular force changes in cells, we used Rho activator (CN01) and ROCK inhibitor (Y27632) to increase or decrease intracellular tension, respectively. CN01 indirectly activates RhoA by suppressing upstream inhibitors, and Y27632 selectively inhibits ROCK. A significant increase in Dsg3 FRET ratio was observed when HaCaT cells were treated with ROCK inhibitor Y27632, while a reduction in FRET ratio was induced by Rho activator CN01. In contrast to the tension sensor, there was no significant Dsg3 FRET ratio change for the tailless control during RhoA activation or inhibition (Figure 2D-F). This data suggests that Dsg3 may be under homeostatic tension and this tension is altered when the global tension within the actin network is modulated.

To study the effect of PV Abs on desmosome tension at cell-cell contact, we treated HaCaT cells expressing Dsg3 tension sensors with AK23 and PVIgG Abs. In the first 30 min of treatment, there were no significant changes in the measured FRET ratio for the four conditions tested (Figure S8). However, after 4h, all conditions resulted in a significantly increased FRET ratio, implying that the tension changes were delayed in response to Ab binding (Figure 2G and H, Figure S9). At this timepoint, compared to controls, FRET ratio increased by around 5% and 10% with AK23 treatments of 2 μg/mL and 10 μg/mL, respectively. PVIgG, which was previously shown to cause more severe junctional dissociation, induced a similar increase in FRET ratio at concentrations of 2 μg/mL and 100 μg/mL as AK23 at 10 μg/mL.

To evaluate the sustained response of Dsg3 tension, we measured the FRET ratio at 8h and 24h after Ab treatments (Figure 2G and H, Figure S9). At 8h, the FRET ratio for AK23 at 2 μg/mL reached its maximum of around 8.5% higher than controls, and subsequently decreased at 24h, indicating that Dsg3 tension loss may start to recover after 24h. This recovery was also observed in AK23 10 μg/mL groups, where the FRET ratio reached a maximum at 4h and began to decrease over 24h. For PVIgG at 2 μg/mL, the FRET ratio was maintained at around 10% higher than controls over the 24h, peaking slightly at 8h. Conversely, for PVIgG at 100 μg/mL, no FRET ratio decrease was observed as treatment went on for 24h. In fact, FRET ratio steadily increased over 24h, suggesting a sustained tension loss with PVIgG treatment. No significant changes in FRET ratio were observed for the tailless control sensor treated with PV Abs, suggesting that the changes in FRET ratio for the tension sensor were the result of changed tension within Dsg3 (Figure S10). Together, our results showed that Dsg3 was sensitive to both intercellular and intracellular tensional changes and that different mechanisms may be at play for AK23 and PVIgG Abs in reducing Dsg3 tension.

### Traction forces at cell-cell adhesions are elevated by PV-associated Abs

After examining the effect of PV Abs on desmosomal disruption and tension loss, we next investigated how this changed tension environment influenced tension at cell-ECM adhesions and cell-cell adhesions as a whole. To do this, we used traction force microscopy (TFM) to quantify the traction stress exerted by PV Ab-treated HaCaT cells (Figure S11). PVIgG induced a small but significant reduction in total traction stress per cell at 4h and 24h, whereas AK23 only reduced it at a concentration of 2 μg/mL at 4h, with no changes at 24h or at either time point with a concentration of 10 μg/mL (Figure 3A and B, Figure S12 and S14A, B). The average traction stress, calculated by dividing the summation of traction stress by the cell area, did not change significantly for any condition, except for a slight increase for PVIgG at 2 μg/mL at 24h (Figure S12, S13A and S14C, D). However, for both PV-associated Abs, we did observe a significant 5- to 10-fold increase in junctional traction stress, the mean of traction stress along the cell-cell contacts (Figure 3C, Figure S15). This supports the idea that intercellular forces maintained by cell-cell junctions may be compromised by PV Ab treatment and cells exert increased grasp on the TFM substrate underneath the cell-cell junctions in response.^21^ It is worth noting that decrease in traction stress per cell from 4h to 24h observation may be influenced by the retraction of keratinocytes induced by PV Abs, marked by the decrease in cell contact area from 4h to 24h (Figure S13B).

**Figure 3.**
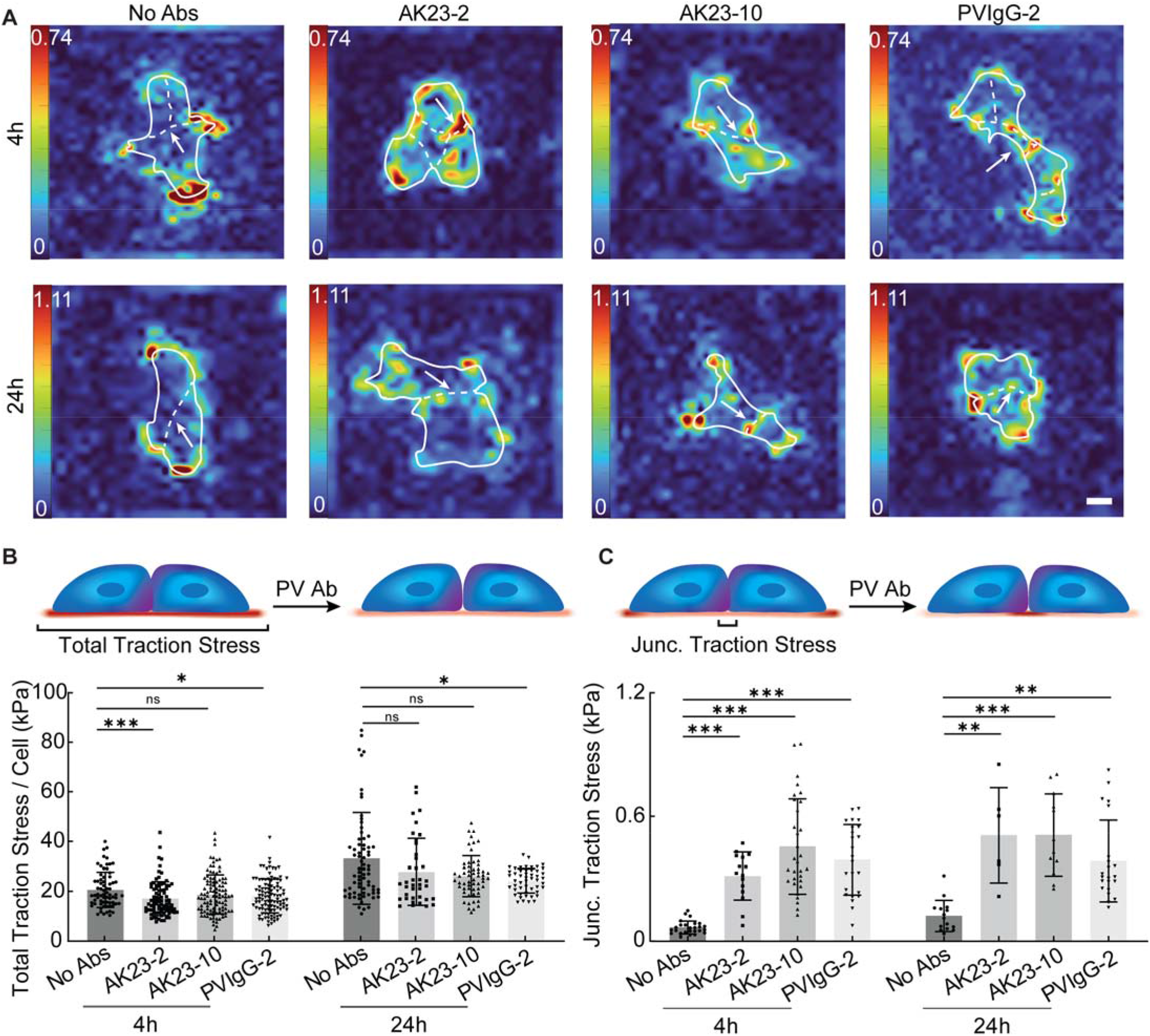
Reduced intercellular tension by PV-related antibodies increases traction stress at the cell-cell contact. (A) Heatmap images of average traction stress at cell-ECM contact treated with 2 µg/mL AK23 (AK23-2), 10 µg/mL AK23 (AK23-10), and 2 µg/mL PVIgG (PVIgG-2) at 4h and 24h. (B) Total traction stress per cell was calculated by dividing the summation of traction stress by the number of cells in the colony. (C) Junctional traction stress was evaluated via the mean traction stress within a defined ROI along cell-cell junctions. All values are mean ± SD (n≥30). * p<0.05, ** p<0.01, *** p<0.005, ns, p>0.05. Scale bar: 15 µm. Three repeats for all conditions.

### PV-associated Ab-mediated tension loss at cell-cell adhesions inhibits RhoA

After revealing the downstream effects of PV Abs on keratinocyte cells within cell-cell and cell-ECM adhesions, we next investigated their potential impact on signaling pathways within the cell. A key regulatory pathway of cell contractility is the Rho pathway, which is controlled by the Rho family of GTPases including RhoA. Therefore, to study the influence of PV Abs on the Rho pathway, we used a RhoA FRET sensor to examine its dynamic activation/inhibition during the time course of PV Ab treatment. Activation of RhoA correlates to an increase in measured FRET ratio, while its inhibition reduces it.^22^ This was first verified by treating HaCaT cells with Rho inhibitor. A decrease in RhoA FRET ratio was observed in the first 30 min of the treatment of Rho inhibitor Y27632 and this inhibition was effective for 24h (Figure 4A and B, Figure S16). With the treatment of AK23 (2 μg/mL), FRET ratio decreased in the first 30 min, similar to Rho inhibitor Y27632 (Figure S16). At 4h, the FRET ratio for the AK23 treatment increased beyond the baseline value as in controls, followed by a significant decrease at 24h (Figure 4A and B). This result was supported by western blot of active RhoA, which showed a significant reduction at the same timepoint (Figure 4C and D). Interestingly, no significant change in total RhoA was observed in all treated groups, except for AK23 treatment for 24h which had a small but significant increase (Figure S17). This suggests that the observed changes in RhoA FRET most likely resulted from Ab-induced conformation changes in RhoA instead of triggering its breakdown or production.

**Figure 4.**
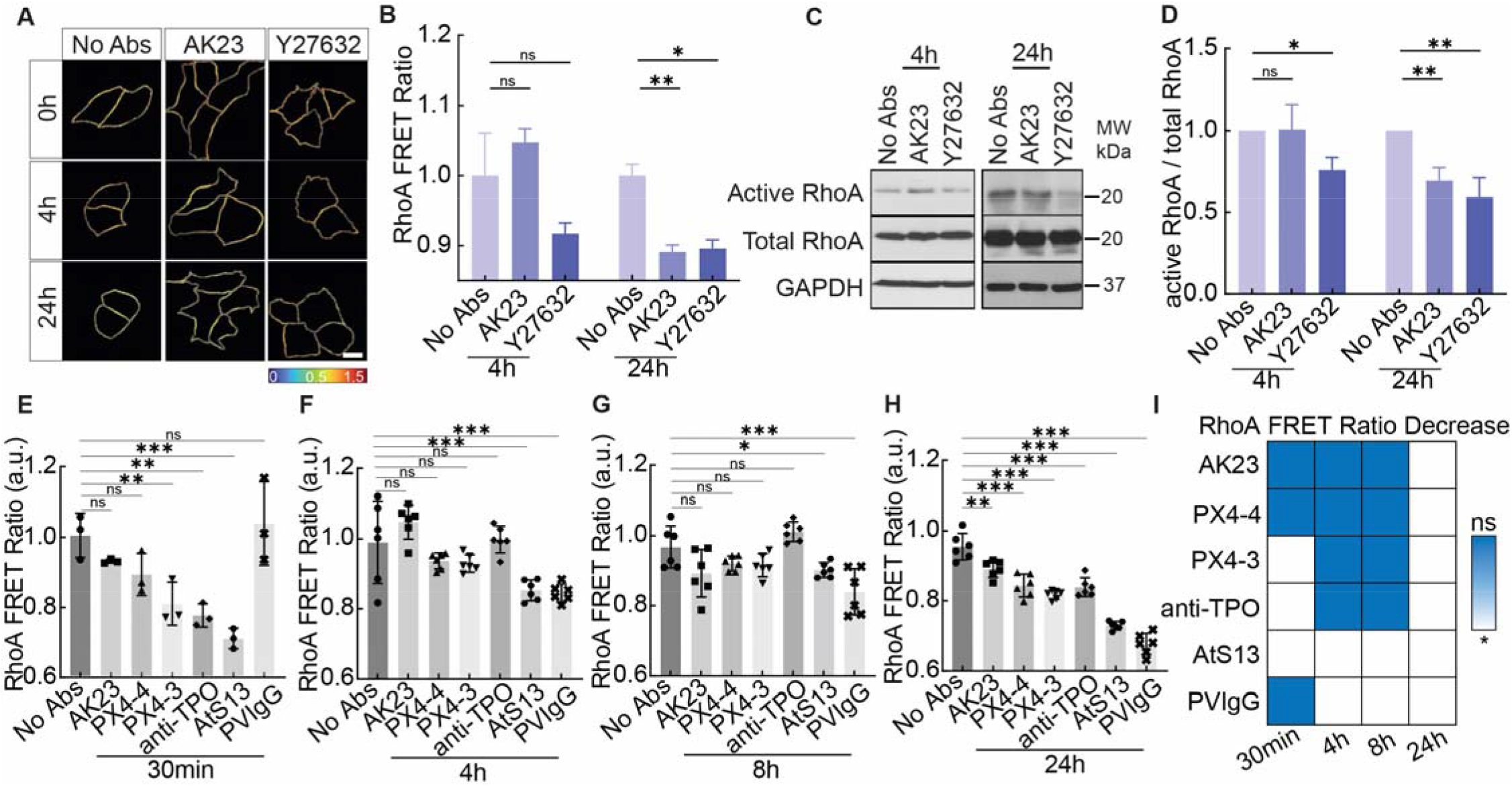
PV Ab-induced tension loss at Dsg3 inhibits RhoA. (A, B) Heatmap images and quantitative analysis of RhoA FRET ratio show the activities of active RhoA in response to 2 µg/mL AK23 and 10 µM Y27632 at 4h and 24h. All values are mean ± SD (n≥30). (C, D) Corresponding western blot results. All values are mean ± SD (n≥3). (E-H) RhoA FRET ratio in HaCaT cells exposed to 2 µg/mL of AK23, PX4-4, PX4-3, anti-TPO, AtS13, and PVIgG at 30 min, 4h, 8h, and 24h. All values are mean ± SD (n≥30). (I) Summary of the significance of RhoA FRET ratio change over 24h for the Abs tested (blue = no change, white = significant decrease). * p<0.05, ** p<0.01, *** p<0.005, ns, p>0.05. Scale bar: 10 µm. Three repeats for all conditions.

After demonstrating that AK23 could alter signaling within the Rho pathway, we next asked if these changes are specific to AK23 or can be extended to other Abs of relevance in PV. Thus, we followed RhoA FRET over 24h under the influence of AK23, PVIgG, and four additional Abs at a concentration of 2 μg/mL (Figure 4E-I, Figure S18), namely anti-thyroid peroxidase (TPO), PX4-3, PX4-4, and AtS13. Anti-TPO is a commercially available Ab directed against human thyroid peroxidase. PX4-4 and PX4-3 are single-chain variable-region fragment (ScFv) anti-Dsg3/Dsg1 Abs isolated by phage display from a patient with mucocutaneous PV and have been described as pathogenic (PX4-3) and mildly to non-pathogenic (PX4-4) in the context of PV.^23^ AtS13 is a PV patient-derived anti-human monoclonal Ab that we developed, which exhibits a 74% heavy-chain homology to anti-TPO Ab and 86% light-chain homology to an anti-desmosome Ab. We have previously demonstrated that similar concentrations of PX4-3 and PX4-4 (10 µg/mL) led to desmosome dissociation,^24^ and our titration experiment also confirmed that 2 µg/mL of PX4-3 is also effective in inducing dissociation (Figure S3B).

The three antibodies with reactivities restricted to Dsg3, namely AK23, PX4-4, and PX4-3 as well as anti-TPO followed a similar pattern of RhoA inhibition over 24h, with no significant effect on RhoA FRET at both 4h and 8h (Figure 4F and G), but a significant reduction at 24h (Figure 4H). However, anti-TPO and PX4-3 also induced a significant early decrease in RhoA FRET ratio at 30 min, while AK23 and PX4-4 did not (Figure 4E). Patient-derived PVIgG, however, lead to a significant and longstanding decrease in RhoA activation at 4h, 8h and 24h (Figure 4E-H), indicating its sustained effect on cell signaling. These results are also supported by the IF retraction and remodeling of actin filaments induced by PV Abs (Figure S19 and S20). Interestingly, the only other Ab showing such a sustained, as well as early, effect was the other fully human, patient-derived Ab AtS13 (Figure 4E-H), which suggests similarities in their effect on cell signaling between these Abs. A summary of the significance of RhoA FRET reduction for all tested Abs at all timepoints is shown in Figure 4I, which demonstrates that all Abs tested, even the presumed non-pathogenic PX4-4, lead to a decrease in RhoA activity at the 24h timepoint, suggesting that all PV-associated Abs ultimately affect cell signaling.

### Tension loss at cell-cell adhesions by PV Abs is restored by RhoA activation

The results of the previous study clearly showed that Rho signaling is altered in PV pathogenesis, as all tested Abs decreased RhoA activity at or before 24h. As the Rho pathway is a key regulator of cell contractility, we next asked if the activation of RhoA could reverse the changes observed in the tension at the different adhesion sites. To investigate this, we measured tension after activating RhoA with Rho activator CN01 in the final 30 minutes of AK23 (2 µg/mL) treatment at 4h and 24h. When added to a 4h treatment of AK23, CN01 did not influence RhoA FRET ratio significantly, but resulted in a nonsignificant increase, similar to AK23 by itself as shown in Figure 4F. However, in contrast, when CN01 was added at the end of a 24h treatment of AK23, it induced a significant increase in RhoA FRET ratio, showing its ability of reversing the effect of AK23 by activating RhoA (Figure 5A). This result was supported by the increase in active RhoA expression, but not total RhoA, with the 24h treatment of AK23 coupled with 30 min of CN01 (Figure 5B and C, Figure S21).

**Figure 5.**
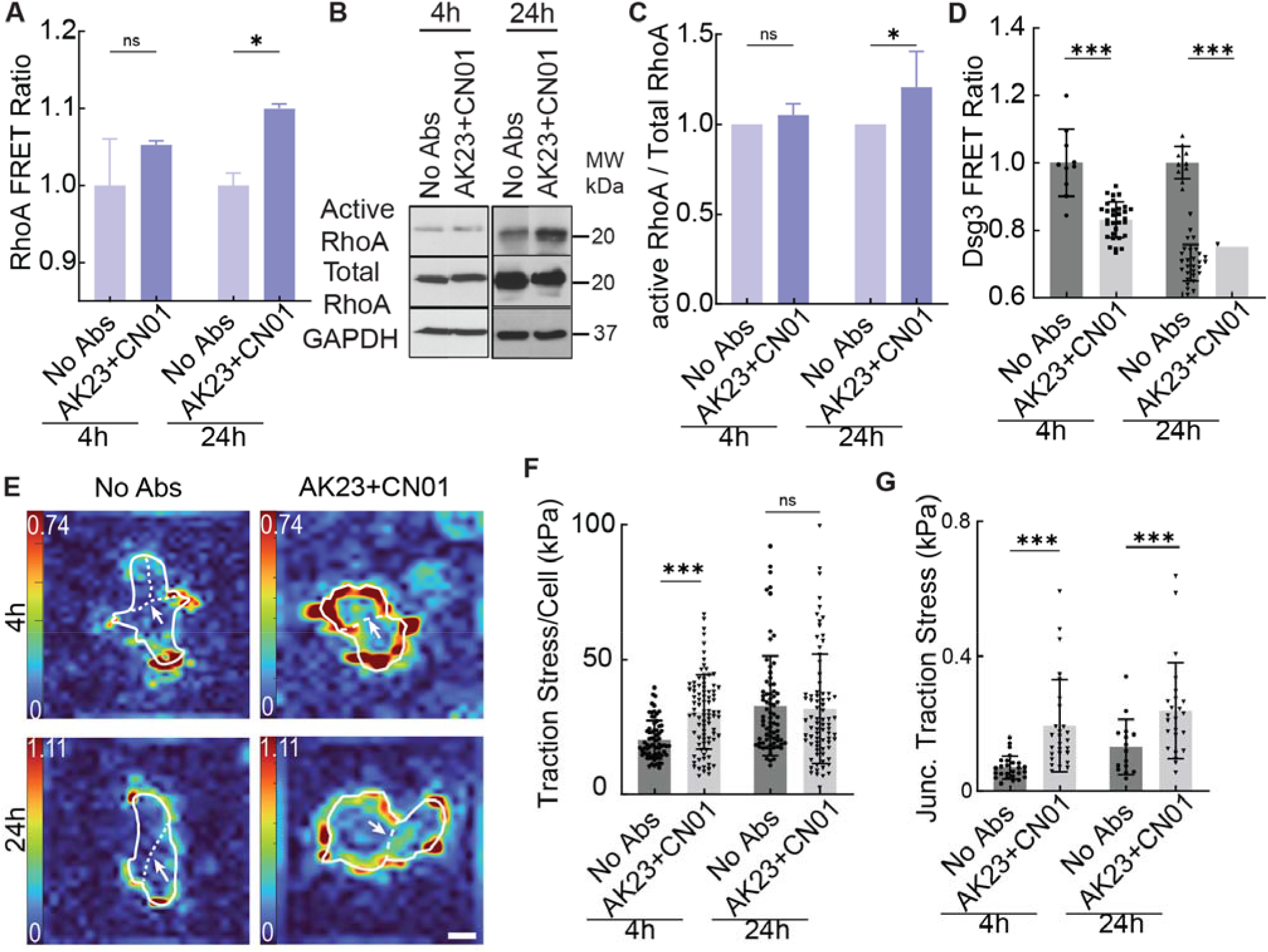
Activation of RhoA restores the tension loss at cell adhesion induced by PV Abs. HaCaT cells were with both 2 µg/mL of AK23 and 1 unit/mL CN01 at 4h and 24h, with CN01 being added in the final 30 min of the treatment. (A) CN01 increases the RhoA FRET ratio at both 4h and 24h. All values are mean ± SD (n≥30). (B, C) Western blot results show the active RhoA expression in HaCaT cells. All values are mean ± SD (n≥3). (D) CN01 reduces the Dsg3 FRET ratio. All values are mean ± SD (n≥30). (E-G) Traction stress and junctional traction stress are increased in HaCaT cells treated with CN01. All values are mean ± SD (n≥30). * p<0.05, ** p<0.01, *** p<0.005, ns, p>0.05. Scale bar: 15 µm. Three repeats for all conditions.

Finally, we investigated how RhoA activation with CN01 could influence the forces experienced at both cell-cell and cell-ECM adhesions. In contrast to treatment with AK23 alone, which increased the FRET ratio measured in Dsg3, CN01 treatment in the final 30 min of both 4h and 24h AK23 treatments reduced the Dsg3 FRET ratio, indicating the restoration of tension across Dsg3 (Figure 5D). CN01 also changed the traction stress exerted by cells. Whereas AK23 treatment alone overall reduced total traction stress per cell slightly, CN01 could increase the total traction stress at 4h and did not significantly change it at 24h (Figure 5E and F). Further, while traction stress beneath the cell-cell junction still increased with additional treatment of CN01, the amount it increased was approximately cut in half compared to treatment of AK23 alone (Figure 5G). We also investigated the dynamics of E-cadherin under the PV Abs and CN01 treatment via fluorescence recovery after photobleaching (FRAP) (Figure S22). We observed that both AK23 (2 μg/mL) and Y27632 reduced the immobile fraction of E-cadherin-GFP while AK23 in combination with CN01 increased it, indicating that Rho inhibition may also lead to a decreasing binding affinity for E-cadherin at cell-cell contacts (Figure S23).

## DISCUSSION

This study investigated the biophysical changes at desmosomes from PV Abs and their effects on RhoA signaling. Using a new FRET sensor on Dsg3, we showed for the first time that it responds to PV Abs with tension loss, which is significant as it provides direct evidence for tension change during Dsg3 disassembly, opening the door for mechanical compensation and mechanotransduction at the cell-cell adhesion as well as cell-ECM adhesions. Specifically, we showed that AK23, a mouse-derived monoclonal anti-human Dsg3 Ab, and PVIgG, the patient serum-derived polyclonal PV Abs, result in different Dsg3 tension loss, junctional dissociation, and cytoskeleton rearrangement. Subsequently, we examined stress at cell-cell contact and cell-ECM adhesions via TFM and observed a reduction in tension at the junction and an increase in local traction stress near the junction, suggesting the compensation of cell-ECM adhesion on cell-cell junctions. These results demonstrate the potential compensation from adherens junctions and focal adhesions during biophysical transformation after PV Ab binding, which has not been previously reported. Further, we showed that these transformations were accompanied by RhoA inhibition at the cell-cell contact and that different levels of inhibition may inform the severity of pathogenicity of different PV Abs. Finally, we demonstrated that stimulating the RhoA pathway could mitigate or reverse these biophysical changes, further implicating the RhoA pathway in PV pathogenesis. These results are significant because they suggest that we could potentially harness cell contractility to restore homeostasis to the cell-cell adhesion as a new type of therapy for PV.

We first demonstrated the dissociation of desmosomal cadherin with PV-associated Abs and the crosstalk between E-cadherin and Dsg3. We observed tightened adherens junctions in keratinocytes with a 4h treatment of 2 µg/mL AK23 while increasing dissociation of E-cadherin was detected in other groups. This implied that the disassembly might be compensated by strengthening adherens junctions via positive feedback between the desmosome-filament and cadherin-actin networks.^25,26^ However, this homeostatic compensation only works when keratinocytes are exposed to a low dosage of mouse monoclonal PV antibodies for a short period. Instead of the countervailing from adherens junctions, vast desmosomal dissociation may induce the activation of EGF receptor, Src, and relative signaling downstream of ligated antigens as well as the elevation of intracellular calcium, and eventually launch the cell death cascades.^27^ We next showed that the binding of PV-associated Abs led to reduced tension across Dsg3 at cell-cell contacts. We observed an increase and decrease in Dsg3 tension induced by Rho activator and inhibitor, respectively, indicating that Dsg3 experiences intracellular tension. Activation of the Rho signaling pathway increases cell contractility, and eventually delivers tension changes to Dsg3 through the desmosome-IF network.^28^ There are recent reports showing that desmosomes are under tension either in homeostasis or under applied load, including findings in desmosomal proteins DP^29^ and Dsg2.^30^ We now know that Dsg3 also experience tension under homeostasis conditions. More importantly, we detected tension loss in Dsg3 after PV Ab treatment, suggesting that Dsg3 may also sense the intercellular tension changes. The binding of PV Abs activates multiple signaling pathways, such as p38MAPK, Src, and Hsp27, which lead to the retraction of IFs from cell-cell contact.^18,31^ IF retraction may trigger crosstalk with actin via plectin and result in the turnover of actin,^32^ which affects cellular tension. Intriguingly, we observed that the tension loss induced by AK23 was slightly recovered at 24h, whereas PVIgG showed no sign of recovery at this timepoint. Further, PVIgG at 100 µg/mL resulted in a continuous tension loss over 24h. This suggests that PVIgG has the capability to induce a more prolonged force imbalance at cell-cell contacts while resisting homeostatic adjustment, which could explain its strong pathogenic effect.

We investigated the interplay between tension across desmosomal cadherins and cell-ECM adhesions and observed that interfering with Dsg3 mediated cell-cell contacts with PVIgG and AK23 changed the distribution of cell-ECM forces underneath the cells. With no Ab treatment, the highest cell-ECM forces were found at the outer edge of small cell colonies. However, the introduction of these Abs elevated cell-ECM forces directly underneath cell-cell contacts inside of these colonies. Compared to the junctional traction stress of untreated cells, both AK23 and PVIgG increased the junctional traction stress at both 4h and 24h. For AK23, the junctional traction stress further increased from a 4h treatment to 24h, whereas for PVIgG the junctional traction stress was not changed from 4h to 24h. These changes in cell-ECM forces could be explained by the result of passive effects of force balancing as tension is lost in the cell-cell junction or could be through an active biochemical response inside the cells. For passive force balancing effects, it would be expected that traction forces beneath cell-cell junctions would increase when tension is lost in cell-cell junctions, and vice versa. However, whereas some tension recovery was observed in Dsg3 when treated with AK23 for 24h using FRET, traction stress continued to increase under cell-cell junctions from 4h to 24h, indicating an active response of the cells to increase traction. Similarly, tension forces within Dsg3 continuously decreased over 24h when treated with PVIgG as revealed by FRET, whereas traction forces underneath cell-cell junctions did not change between 4h and 24h, again suggesting an active response by the cells to modulate traction. These observations support the claim that there is crosstalk between cell-cell and cell-ECM adhesions to help balance forces on an individual cell, which has been observed previously.^21^

To further understand the signaling pathways induced by PV Abs, we investigated the activation of RhoA using a RhoA FRET sensor. The inhibition of the active RhoA was visualized by the reduction of FRET ratio in keratinocytes exposed to ROCK inhibitor. Interestingly, in cells treated with 2 µg/mL AK23, even though changes were not statistically significant, we found that RhoA activity was inhibited at 30 min, activated at 4h, and again inhibited at 24h. This indicates that RhoA signaling pathway is sensitive and time dependent.^33,34^ The transient activation of RhoA may result from the coordination of different cytoskeletal networks. Specifically, crosstalk between actin and IFs increases forces at E-cadherin and triggers cadherin signaling, which has been shown to inhibit the activation of RhoA through p190RhoGAP.^35^ Further, our data support the idea that tension loss within Dsg3, which occurs within 4h of exposure to PV Abs, is upstream of changes in RhoA signaling, which did not significantly change until 24h, at least for AK23.^36,37^

Interestingly, the anti-Dsg3 Ab AK23 and anti-Dsg3-dominant PVIgG are not the only Ab sources found to induce RhoA activation in our study. Other Abs with relevance to PV pathogenesis induce RhoA activation as well, however with slightly different timing and duration. While all Abs used in this study ultimately activate RhoA at the 24h timepoint, only patient derived PVIgG and fully human patient derived Ab AtS13 led to a significant sustained decrease in RhoA activation at all timepoints (with the exception of 30 min for PVIgG only), suggesting similarities in their effect on cell signaling between these Abs. Of note, the scFv anti-Dsg3 Ab PX4-4, thought to be mildly- or non-pathogenic, did not activate RhoA at 30 min, while PX4-3, thought to be pathogenic, did activate RhoA at this early timepoint. It remains to be seen if this difference in RhoA activation can account for the differences in pathogenic potential of these Abs. A monoclonal anti-TPO Ab was identical to PX4-3 in terms of RhoA activation timing, both without the sustained activation seen by the patient-derived Abs, though.

We also evaluated the dynamics of E-cadherin, tension in Dsg3, and traction force in focal adhesions after activating RhoA, and found that RhoA activation provides positive feedback on AK23 induced disruption, indicating that RhoA has the effect of stabilizing cell-cell contacts.^28,38–40^ This benefit could arise from multiple functions of RhoA in cells. First, the active RhoA directly contracts actomyosin which promotes the rearrangement of actin and then launches the engagement and clustering of E-cadherin at cell-cell contact.^41–43^ In addition, it phosphorylates adducin, a protein organizing the cortical actin cytoskeleton, which protects desmosomal cohesion in keratinocytes and modulates desmosomal turnover and plasticity. The loss of the actin-binding protein α-adducin results in reduced desmosome numbers and prevents the cohesion of desmosomal molecules in cultured keratinocytes.^44^ Moreover, RhoA activation regulates the spatial organization of traction stress and the direction of cell motion.^45^ Finally, RhoA activation stimulates mDia1-mediated F-actin assembly at junctions and stabilizes E-cadherin at multicellular vertices.^28,46,47^

The results of this study help illuminate the signaling pathways initiated by PV-associated Abs and demonstrate crosstalk between cell-cell and cell-ECM adhesions (illustrated in Figure S24). When PV Abs attack the intercellular domain of Dsg, the desmosome linkage is disrupted, leading to the release in tension across the desmosome and eventual retraction of IFs. The compromised tension state leads to signaling that increases cell-ECM forces, potentially mediated by RhoA, which could be the result of increased interaction between focal adhesions and actin filaments or clustering of focal adhesions. Stimulation of RhoA can reverse some of the negative effects induced by PV Abs, including mitigating the rise of junctional traction stresses and restoring tension within Dsg3. Taken together, the results of this study point to cell contractility as a potential therapeutic target to stabilize the mechanical environment of keratinocytes and combat the pathological effects of PV. Further development of drugs that target biomechanical signaling pathways including RhoA have the potential to provide therapy with minimal side effects compared to traditionally used immunosuppression strategies.

### Limitations of the study

Several caveats need to be taken into consideration. We showed that desmosomal cadherin tension loss can be induced by PV antibodies and that RhoA inhibition at the cell-cell contact may potentially regulate the tension loss at the desmosome, since restoring RhoA ameliorated some of the tension loss. However, whether restored tension in Dsg3 is the result of enhancing the interaction between intercellular domains or else through tension transmitted by the cell membrane remains to be explored. Potential mechanisms that govern the observed tension restoration, including the protective pathways in PV such as p38 MAPK, still need to be explored. In addition, our results showed PV Ab induced tension loss at Dsg3 and the crosstalk between desmosomes and adherens junctions. However, whether PV Ab binding also results in tension loss at E-cadherin is not covered. FRET tension sensors engineered into E-cadherin, and their co-expression with Dsg3 tension sensors may provide a full account of the tensional dynamics at the cell-cell adhesion during PV Ab induced biophysical response. Meanwhile, though we evaluated retraction of IFs and discussed the potential relationship between Dsg3 tension loss and cytoskeletal remodeling, how force transfers from the cell-cell junction to IFs and whether IFs load the tension changes remain unclear. Furthermore, the FRET tension sensor has its own limitations. It can only measure force changes within nanonewtons and is not compatible with all cell types. Lastly, serum-purified PV Abs vary from patient to patient. Our data showed the inhibition effect of PV Abs on RhoA activation. It remains to be examined whether the effects are shared across different patient serum-sourced PV Abs.

## ACKNOWLEDGMENTS

This study was supported by fundings from NSF (1826135, 2143997) (R.Y.), NIH R35GM150623 (R.Y.), NIH P20GM113126 (R.Y.), NIGMS P20GM104320 (J.Y.L.), NIAMS R15AR072959 (J.K.W.), NIGMS R35GM119617 (D.E.C.) and NIAMS AR068096 (D.E.C.). G.M. and J.R. are supported by the NSF Graduate Research Fellowship (2034837). We thank Professor Amy Payne from University of Pennsylvania for sharing the pathogenic and nonpathogenic antibodies (PX4-3 and PX4-4) and Jason Manning from Cornell University for help in the anisotropic analysis.

## AUTHOR CONTRIBUTIONS

R.Y., A.A.S. and D.E.C. conceived and designed research; X.J., J.R., E.K., K.S., M.L., G.M., Y.H., B.S., H.Z. performed experiments; X.J., J.R., E.K., K.S., M.L., Y.H., H.Z., J.Y.L., C.H., F.M. analyzed data. J.K.W. introduced RhoA FRET sensors to keratinocytes; S.R.B. and D.E.C. constructed and characterized the Dsg3 FRET sensor. K.S., M.L. and A.A.S. developed the patient derived antibodies. All authors contributed to discussion and approved final version of the manuscript.

## STAR METHODS

### KEY RESOURCES TABLE

| REAGENT or RESOURCE | SOURCE | IDENTIFIER |
| --- | --- | --- |
| **Antibodies** |  |  |
| Dsg3 antibody | Abcam | Cat # ab231309; RRID: N/A |
| E-cadherin antibody | Abcam | Cat # ab231303; RRID: AB_2923285 |
| Alexa Fluor 647 | Thermo Fisher | Cat # A-21235; RRID: N/A |
| Alexa Fluor 488 | Thermo Fisher | Cat # A-11001; RRID: N/A |
| Alexa Fluor 488 Phalloidin | Thermo Fisher | Cat # A12379; RRID: N/A |
| Anti-Dsg3 antibody (AK23) | MBL | Cat # D218-3; RRID: AB_591236 |
| PX4-4 | This study | N/A |
| PX4-3 | This study | N/A |
| Anti-TPO | Novus Biologicals | Cat # NBP1-50811; RRID: N/A |
| AtS13 | This study | N/A |
| PVlgG | This study | N/A |
| rabbit anti-GFP | Abcam | Cat # ab6556; RRID: AB_305564 |
| 6 nm colloidal gold conjugated goat-anti-rabbit IgG | Electron Microscopy Sciences | Cat # 25103; RRID: N/A |
| rabbit anti-GFP | Cell Signaling | Cat # 2956S; RRID: N/A |
| mouse anti-desmoplakin | Fitzgerald | Cat # 10R-D108a; RRID: AB_1284113 |
| mouse anti-E-cadherin | BD Biosciences | Cat # 610181; RRID: AB_397580 |
| anti-active RhoA GTP | NewEast Biosciences | Cat # 26904; RRID: AB_1961799 |
| EGTA | Biotechne | Cat # 2807; RRID: N/A |
| rabbit anti-RhoA | Cell Signaling | Cat # 2117S; RRID: N/A |
| mouse anti-GAPDH | Santa Cruz Biotechnology | Cat # SC-32233; RRID: AB_627679 |
| HRP-linked anti-mouse IgG | Jackson ImmunoResearch | Cat # 115-035-166; RRID: AB_2338511 |
| HRP-linked anti-rabbit IgG | Jackson ImmunoResearch | Cat # 111-035-003; RRID: AB_2313567 |
| **Chemicals, peptides, and recombinant proteins** |  |  |
| Dsg3 tension sensor | This study | N/A |
| Dsg3 tailless control sensor | This study | N/A |
| pTriEx-RhoA FRET WT biosensor | Addgene | Plasmid #12150; RRID: Addgene_12150 |
| LZBob-neo-vector | Gift from James K. Wahl, III | N/A |
| ROCK Inhibitor (Y27632) | Sigma-Aldrich | Cat # SCM075 |
| Y27632 | Selleck Chemical LLC | Cat # S1049 |
| Rho activator | Cytoskeleton | Cat # CN01-A |
| Polybrene | Sigma | Cat # 28728-55-4 |
| G418 | Invivogen | Cat # ant-gn-1 |
| Aurion Blocking Solution | Electron Microscopy Sciences | Cat # 25595 |
| BSA | Thermo Fisher Scientific | Cat # 37520 |
| glycine | Millipore Sigma | Cat # 410225 |
| Tween 20 | Millipore Sigma | Cat # P7949 |
| protease-inhibitor cocktail | Sigma Aldrich | Cat # S8830 |
| protein assay dye reagent | Bio-Rad | Cat # 5000006 |
| ECL chemiluminescence | Santa Cruz Biotechnology | Cat # sc-2048 |
| **Critical commercial assays** |  |  |
| pAdEasy adenoviral vector system | Agilent | Cat # 240009 |
| MBL (MBL2) Human ELISA Kit | ThermoFisher | Cat # EHMBL2 |
| Zeba Spin Desalting Columns | ThermoFisher | Cat # 89882 |
| Pierce Centrifuge column | ThermoFisher | Cat # 89896 |
| Melon Gel IgG Spin Purification Kit | ThermoFisher | Cat # 45206 |
| Gel with 0.2 μm fluorescent beads | Matrigen | Softview 35/20 Glass |
| **Experimental Models: Cell Lines** |  |  |
| HaCaT cells | Gift from James K. Wahl, III | N/A |
| 293A cells | ThermoFisher | Cat # R70507 |
| Phoenix cells | Gift from James K. Wahl, III | N/A |
| **Recombinant DNA** |  |  |
| E-cadherin-GFP | Addgene | Plasmid # 28009; RRID: Addgene_28009 |
| **Software and algorithms** |  |  |
| MATLAB R2020b | MathWorks | https://www.mathworks.com/products/new_products/release2020b.html |
| Fiji | NIH | https://imagej.net/software/fiji/ |
| GraphPad Prism | GraphPad Prism 9 | https://www.graphpad.com/updates/prism-900-release-notes |
| Zen | Zeiss | https://www.micro-shop.zeiss.com/en/de/softwarefinder/software-categories/zen-blue/zen-blue-pro |
| LASX Office | Leica | https://www.leica-microsystems.com/products/microscope-software/p/leica-las-x-ls/downloads/ |
| ImageLab Software | Bio-Rad | https://www.bio-rad.com/en-us/sku/12012931-image-lab-software-usb-drive?ID=12012931 |
| **Other** |  |  |
| Confocal microscope | Zeiss | LSM800 |
| Inverted Fluorescent microscope | Leica | DMI8 |
| Leica EM UC6i ultramicrotome | Leica | EM UC6i |
| Transmission electron microscope | Jeol | JEM-1230 |
| 3D Printer | Stratasys | Objet500 Connex3 |
| Nanodrop® spectrophotometer | Thermo Scientific | Cat # ND2000CLAPTOP |
| ChemiDoc imager | Bio-Rad | https://www.bio-rad.com/en-us/category/chemidoc-imaging-systems?ID=NINJ0Z15 |
| PTFE resin molds | Ted Pella, Inc. | Cat # 10509 |
| Aclar film | Ted Pella, Inc. | Cat # 10501 |
| polyvinylidene fluoride (PVDF) membranes | Bio-Rad | Cat # 1620177 |

### RESOURCE AVAILABILITY

#### Lead contact

Further information and requests for resource and reagents should be directed to and will be fulfilled by the Lead Contact, Ruiguo Yang.

#### Materials availability

This study did not generate new unique reagents.

#### Data and code availability

All data reported in this paper will be shared by the lead contact upon request. This paper does not report original code. Any additional information required to reanalyze the data reported in this paper is available from the lead contact upon request.

## METHOD DETAILS

### Constructs

Tension changes were evaluated in keratinocytes (HaCaT) expressing Dsg3 FRET sensors composed of mouse sourced Dsg3 cDNA and tension sensor module (TSMod), where mTFP1 and Venus were connected by an amino acid elastic linker. The Dsg3 tension sensor was created by ligating TSMod to the mouse Dsg3 cDNA with modified SalI and NotI sites between intracellular anchor and intracellular catenin-binding site. The Dsg3 tailless control sensor was built by removing the DP connecting cytoplasmic tail of Dsg3 cDNA. Reconstructed Dsg3 FRET sensors were engineered into the adenoviral Dsg3 FRET constructs via pAdEasy Adenoviral Vector System. The constructs were then transformed into 293a cells and adenoviral conditioned media was collected based on a previously published protocol for transfection^48^. Antibody binding affinity of mouse sourced Dsg3 has been proved using passive transfer mouse model in pemphigus studies.^49,50^

The RhoA sensor construct was made by recombining commercial RhoA sensor plasmid from Addgene into the LZBob-neo-vector, which is a modified LZRS-ms neo-vector with multiple cloning sites for increasing cDNA fragment cloning. pTriEx-RhoA FLARE.sc WT biosensor (Addgene, Plasmid # 12150) was digested and ligated to LZBob-neo-vector to make retroviral RhoA sensor construct. E-cadherin–green fluorescent protein (E-cad–GFP) construct was generated via the LZBob-neo-vector. Human E-cadherin cDNA was digested and ligated to the vector from retroviral E-cad-GFP fusing protein construct. The retroviral constructs were transformed into Phoenix cells for packaging and amplifying. Phoenix cells were then cultured in calcium-dependent media which was made from Dulbecco’s modified Eagle’s medium (DMEM) (11965092, Thermo Fisher Scientific, calcium concentration: 1.7mM) supplemented with 10% fetal bovine serum (FBS) (A3382001, ThermoFisher Scientific), 1% penicillin-streptomycin (15140122, ThermoFisher Scientific), and 1% GlutaMAX for more than 2 days. The viral conditioned culture medium was collected and filtered with a 0.45 μm syringe filter for transfection.

### Cell culture

The immortalized human keratinocytes (HaCaT cells) were cultured in low calcium media made from no-calcium DMEM (21068028, ThermoFisher Scientific), 10% FBS and 1% penicillin-streptomycin at 37°C supplied with 5% CO_2_ to maintain the dividing state.

### Cell transfection

The adenoviral-based transfection was processed ahead of each experiment. HaCaT cells with around 50% confluency were infected by culturing in the adenoviral conditioned media with 4 μg/mL polybrene (28728-55-4, Sigma) for 7h at 37°C. For retroviral-based transfection, HaCaT cells were infected by culturing in the retroviral conditioned media with 4 μg/mL polybrene for 7h at 32°C and then sorted by culturing in low calcium media with 500 μg/mL G418. Cells expressing the FRET sensor were further selected via flow cytometry. Cell viability was confirmed using LIVE/DEAD™ Viability/Cytotoxicity Kit (Thermofisher, L-3224) after transfection (Figure S25). All infected HaCaT cells were cultured in low calcium media at 37°C supplied with 5% CO_2_ and were then changed into calcium-dependent media overnight ahead of drug treatments and imaging.

### Purification of PVIgG

PVIgG was purified from the biopsy-confirmed PV patient serum that had been determined to be anti-Dsg3 positive (135.99 U/mL) and anti-Dsg1 negative (7.96 U/mL) by ELISA (MBL). Purification of PVIgG was performed using Zeba column (ThermoFisher), Pierce Centrifuge column and Melon Gel IgG Spin Purification Kit (ThermoFisher) according to manufacturer’s instructions. The protein concentration of purified fractions was analyzed by a Nanodrop® spectrophotometer. Fractions were no longer collected when protein values fell below 0.2 mg/mL. To determine PVIgG purity, a sample of each of the highest yielding fractions was separated by SDS-PAGE gel stained by Coomassie. Bands were imaged on a ChemiDoc imager and analyzed using ImageLab Software (Bio-Rad). Purity of PVIgG in a fraction was expressed as percent of IgG (H+L chains) out of the total protein in that fraction’s sample. Typical yields ranged from 0.3 – 2.3 mg/mL (70-88% purity).

### Antibody treatment

HaCaT cells were grown in no calcium media to reach 80% confluency and changed into calcium-dependent media overnight before the drug treatment. All drug treatments were performed in calcium-dependent media. PV Abs including PX4-4, PX4-3, anti-thyroid peroxidase (anti-TPO) (NBP1-50811, Novus Biologicals), AtS13, and PVIgG were used at a concentration of 2 µg/mL. PX4-4 and PX4-3 are derived from the single-chain variable-region fragment (ScFv) monoclonal Abs isolated from a patient with mucocutaneous PV (kindly provided by Dr. Aimee Payne, University of Pennsylvania), and AtS13 is a PV patient-derived anti-human monoclonal Ab, which exhibits a 74% heavy-chain homology to anti-TPO Ab and 86% light-chain homology to an anti-desmosome Ab (developed by the A. A. Sinha group, University at Buffalo). PVIgG is polyclonal Abs purified from a Dsg3-containing patient serum (developed by the A. A. Sinha group, University at Buffalo). Anti-Dsg3 antibody (AK23) (D219-3, MBL) was used at a concentration of 2 μg/mL or 10 μg/mL based on the experiments. EGTA was used at a concentration of 5 mM. Rho activator I (CN01) (Cytoskeleton) was used at a concentration of 1 unit/mL. For Dsg3 tension and tailless sensor, and for RhoA sensors, HaCaT cells were treated with CN01 for 30 min before imaging. ROCK inhibitor (Y-27632) (SCM075, Sigma) was used at a concentration of 10 μM for 12h.

### Traction force microscopy

HaCaT cells were seeded on collagen-coated gel with a stiffness of 12 kPa. The surface of the gel was embedded with 0.2 μm fluorescent beads (Ex/Em: 505/515) at a 1:800 bead dilution ratio (Softview 35/20 Glass, Matrigen). Fluorescent beads with and without cell colonies were imaged using Leica DMi8 inverted microscope (Leica) with a 20X/0.4NA objective. Traction stresses were calculated via the particle image velocimetry (PIV) plugin, and the Fourier transform traction cytometry (FTTC) plugin in ImageJ (NIH). The area of cell colonies was quantified and measured using the polygon tool in ImageJ. Junctional traction stress was calculated with a custom-written MATLAB code (Figure S11, Figure S15). More than 30 images were collected from different fields of view for each sample and at least three independent samples were analyzed for each group.

### Immunogold electron microscopy

HaCaT cells expressing the desmoglein-3 sensor were grown on Thermanox coverslips and fixed with 4% paraformaldehyde/0.1% glutaraldehyde in 0.1M Millonig’s buffer for 1h. Following fixation, samples were washed briefly (3 × 5 min) in phosphate buffered saline (PBS). The samples were then dehydrated through a graded series of ethanol (30 to 100%, 10 min at each step). Following dehydration, samples were infiltrated with 3:1 100% ethanol:LR White (1h on a rotator), 1:1 100% ethanol:LR White (1h on a rotator), and 1:3 100% ethanol:LR White (2h on a rotator). The samples were then infiltrated overnight in LR White at 4°C. The following day, the samples were flat embedded (cell side up) in polytetrafluoroethylene (PTFE) resin molds (Ted Pella, Inc., Redding, CA, USA), overfilled with LR White, and sealed with Aclar film (to avoid any exposure of the LR White to oxygen during polymerization). Polymerization of LR White was done either in an oven (set to 55°C) or under UV light (in a chest over dry ice) for 24h. 60 nm sections were cut (Leica EM UC6i ultramicrotome, Wetzlar, Germany) and collected on 300 mesh formvar-coated nickel grids. Prior to immunolabelling, grids containing sections were incubated (face down) on 100 μL drops of Aurion Blocking Solution (Electron Microscopy Sciences, Hatfield, PA, USA) for 1h at room temperature. The grids were then washed (face down) on 100 μL drops of PBS (with 0.1% BSA) on a rotary stir plate, set to the lowest setting (3 × 5 min). The grids were incubated (face down) on 150 μL drops of rabbit anti-GFP (Abcam, Cambridge, MA, USA, ab6556), diluted 1:50 in PBS (with 0.1% BSA), overnight in a sealed humidity chamber at 4°C. The following day, the grids were washed (face down) on 200 μL drops of PBS (3 × 5 min, a rotary stir plate, set to the lowest setting). The grids were then incubated (face down) on 150 μL drops of 6 nm colloidal gold conjugated goat-anti-rabbit IgG (Electron Microscopy Sciences), diluted 1:20 in PBS (with 0.1% BSA), for 2h in a sealed humidity chamber at room temperature. Following incubation in the secondary antibody, the grids were washed in PBS, with 0.1% BSA (3 × 5 min), PBS (3 × 5 min), and then distilled water (5 min). Prior to imaging, the grids were stained with 2% aqueous uranyl acetate (10 s) and Reynold’s lead citrate (10 s). The grids were imaged with a Jeol JEM-1230 transmission electron microscope (TEM) equipped with a Gatan Orius SC1000 CCD camera. At least three independent samples were analyzed for each group.

### Protein extraction and Western blot analysis

Cells transfected with the Dsg3 tension sensor, tailless control, as well as cells that were not transfected were scraped in TNE buffer (50 mM Tris acetate pH 8.0, 1% Triton X-100, 2 mM EDTA, PMSF) and shaken on ice for 15 min. The extract was mixed vigorously with a pipet for 1 min and then centrifuged at 14000xg for 15 min. The supernatant was kept as the soluble fraction, and the insoluble pellet was washed with TNE buffer and centrifuged again for 5 min. The pellet was suspended in sample buffer and briefly sonicated to solubilize the pellet. Samples were ran with SDS-PAGE and stained with anti-GFP (Cell Signaling Technology) or anti-Dsg3 (produced using a previously reported protocol antibodies to detect the FRET sensor and endogenous Dsg3, respectively.^51^ Analysis of the blots was done using a LI-COR Odyssey imager.

### Immunostaining

Drug-treated HaCaT monolayers were washed with phosphate buffer saline (PBS) (Thermo Fisher) three times and fixed with 4% paraformaldehyde for 10 min for fixing Dsg3, E-cad and F-actin, and ice-cold methanol-acetone (1:1) for 5 min for intermediate filaments. After fixation, monolayers were permeabilization with 0.1% Triton X-100 for 5 min. Next, monolayers were blocked for 1h using 1wt% BSA (37520, Thermo Fisher Scientific), 22.52 mg/mL glycine (410225, Millipore Sigma) in DPBST (DPBS+ 0.1% Tween 20 (P7949, Millipore Sigma)) and were washed with PBS three times for 5 min each before primary and secondary treatments. Dsg3 antibodies (1:200, Abcam), E-cad antibodies (1:200, Cell Signaling Technologies), Alexa Fluor 647 (1:100, Thermo Fisher) and Alexa Fluor 488 (1:100, Thermo Fisher) were used to stain Dsg3, E-cad and intermediate filaments step by step, with each staining step lasting for 1h. F-actin was stained with Alexa Fluor 488 Phalloidin (1:100, Thermo Fisher) for 30 min. HaCaT monolayers after staining were washed with PBS three times for 5 min each and then imaged. For co-staining of Dsg3 and DP, HaCaT monolayers were fixed in ice cold methanol for 5 minutes. Next, cells were blocked using 5wt% BSA for 1h. Cells were stained with rabbit anti-GFP (Cell Signaling, 2956S) to re-label the DSG-3 sensor after methanol fixation (GFP antibodies react against Venus in the sensor), and mouse anti-desmoplakin (Fitzgerald, 10R-D108a), followed by anti-rabbit Alexa Fluor 488 and anti-mouse Alexa Fluor 568. HaCaT monolayers after staining were washed with PBS three times for 5 min each and then imaged.

### Live-cell imaging

HaCaT monolayers were cultured in 29 mm glass bottom dish (D29-20-1.5H, Cellvis) and were imaged with a confocal microscope (LSM800, Zeiss) equipped with an incubation system for time-lapse observation. A dish cap with influx and outflux channels was designed via SolidWorks and printed using an Objet500 Connex3 PolyJet 3D Printer with VeroClear (Stratasys) for changing into drug-containing media conveniently. All 0 min images were captured before drug treatments.

For FRET experiments, images were captured from both donor and acceptor channels after stimulating the donor. To observe the response of cells after drug treatments in real time, a timelapse module was used to automatically capture images every 5 min with a total observation of 30 min in the same field of view. In addition, monolayers with drug treatments were also imaged at 4h, 8h and 24h. At least 15 images from different fields of view were taken at each time point. HaCaT cells with drug-containing media were kept in the incubator between imaging time points. Images were captured using a 63X/1.4NA oil immersion objective with suitable excitation and emission settings of laser and filter.

For FRAP experiments, a bleaching module was used for automatically processing laser bleaching and capturing pre- and post-bleaching images. Regions of interest, including bleached, unbleached, and background, were selected during the imaging. Images were captured every 4 seconds for a total of 200 seconds. At least 5 pre-bleaching images were captured for each sample. Imaging was processed using a 40x/1.3NA oil immersion objective with suitable excitation and emission settings of laser and filter. Imaging parameters, including image size, laser power, master gain, and objective pinhole diameter, were optimized for each protein and kept consistent between groups.

### Immunoprecipitation and immunoblotting

HaCaT cells were seeded on 6-well plates and incubated with AK23 mAb (2 or 10 µg/mL), PVIgG (2 or 100 µg/mL) or Y27632 (ROCK inhibitor; Sellect Chemical LLC, 10µM) for 4h or 24h. CN01 (RhoA activator, 1 unit/mL) was incubated for 20 min due to the toxicity. HaCaT cells were lysed with 1% Triton X-100 buffer (50 mM Tris–HCl, pH 7.4, 150 mM NaCl, 5 mM EDTA, 2 mM dithiothreitol, 1 mM PMSF and 1% Triton X-100) with a protease-inhibitor cocktail (S8830; Sigma Aldrich) and phosphatase inhibitor (Sigma Aldrich). Whole-cell lysates were incubated on ice for 30 min and then centrifuged at 14000g for 20 min at 4°C. The supernatants were collected and quantified with protein assay dye reagent (Bio-Rad), then used for western blot. For immunoprecipitation of active RhoA, the supernatants were incubated overnight with 1 µg/mL anti-active RhoA GTP (NewEast Biosciences), followed by incubation with protein A/G PLUS-agarose for 1h. For western blotting, the samples were separated by 8% or 12% SDS-PAGE and transferred to polyvinylidene fluoride (PVDF) membranes (Bio-Rad). After blocking with 5% skim milk, the blots were incubated with primary antibody, mouse anti-E-cadherin (BD Biosciences; 610181), mouse anti-Dsg3 (Abcam), rabbit anti-RhoA (2117S; Cell Signaling Technology) or mouse anti-GAPDH (SC-32233; Santa Cruz Biotechnology) at 4°C overnight. After three times washing with 1X TBST, the blots were incubated with the HRP-linked anti-mouse or anti-rabbit IgG (Jackson Immunoresearch) at room temperature for 40 min, followed by washing with 1X TBST 3 times. Immunoreactivity of protein was detected by ECL chemiluminescence (Santa Cruz Biotechnology). At least three independent samples were analyzed for each group.

### Image processing and quantification analysis

To minimize the effect of background noise and non-real signals on analysis, rolling ball background subtraction was performed on all raw images via ImageJ before running the image analysis. Pixels with saturated fluorescent intensity were also removed before analysis. All normalized data was calculated by dividing the data of the drug-treated group by that of the control groups.

#### FRET analysis

A binary mask was generated based on the donor image using the drawing tool in ImageJ and multiplied to both donor and acceptor images to outline the cell-cell junctions. FRET ratio was calculated by dividing the acceptor images by donor images and averaging all pixels. Pixels with overflow FRET ratio were removed from both images and the FRET ratio was recalculated based on all sorted pixels. More than 30 images were collected from different fields of view for each sample and at least three independent samples were analyzed for each group. The FRET ratio of experimental groups was then calculated by averaging FRET ratio values from each image in each group. Ratiometric images were generated by dividing the masked and corrected acceptor images by corresponding donor images via MATLAB (Figure S6).

#### Dsg3 full width at half maximum analysis

Distribution curves were plotted by the fluorescent intensity crossing the cell-cell junction at each pixel position. Specifically, a line was drawn from one cell nucleus to another. Profile of the line with Dsg3 intensity and spatial information was plotted and shown as the distribution curve (Figure S1). The initial point of the line was set as 0 and the total length of the line was determined via the total pixel number. Full width at half maximum (FWHM) was calculated based on the spatial information at half of the normalized peak intensity. The normalized peak intensity was calculated by dividing the peak intensity by the mean intensity from 5 pixels at each end of each line to control for background noise. A normal distribution curve was usually seen on the plot with spatial information on the X-axis and intensity value on the Y-axis. Half of the normalized peak intensity corresponded to two values on the X-axis. The difference between the two X values was used to represent FWHM for quantifying spatial information of cell-cell junctions (Figure S1). At least 20 lines were drawn across junctions for one image and the FWHM of the image was calculated by averaging the FWHM value of each line. For each sample, more than 10 images were captured from different fields of view. At least three independent samples were processed for each group. The final FWHM of a group was calculated via averaging the FWHM value of each image.

#### FRAP analysis

FRAP data in excel format was exported via ZEN software and processed using the easyFRAP web-based tool and GraphPad Prism 9 ^52^. Specifically, values were corrected by the background and photofading via the values of the background region and unbleached region. Abnormal FRAP data after the correction was removed before the fitting analysis. Corrected FRAP was then fitted by the nonlinear fitting model built in Prism 9 and the mobile fraction was directly extracted from the fitting result. The immobile fraction was calculated by subtracting the mobile fraction from 1. At least 5 separate FRAP data were captured for each sample and 3 repeats were performed for each treated group. FRAP curves of each group were normalized and averaged to generate one representative FRAP curve.

## QUANTIFICATION AND STATISTICAL ANALYSIS

Statistical data was analyzed using GraphPad Prism 9. All results were presented as mean ± standard deviation. Quantile-quantile (Q-Q) plot was used for the normality test. One-way ANOVA with Tukey multiple comparisons test was performed to compare the significance among groups. To compare the difference between several different time points with three or more groups, two-way ANOVA with Tukey multiple comparisons test was used. A p-value of less than 0.05 was considered statistically significant.

