## Supplemental Information for "Desmosomal Cadherin Tension Loss in Pemphigus Vulgaris Mediated by the Inhibition of Active RhoA at Cell-Cell Adhesions"

**This PDF file includes:**

Figures S1 to S25

SI References

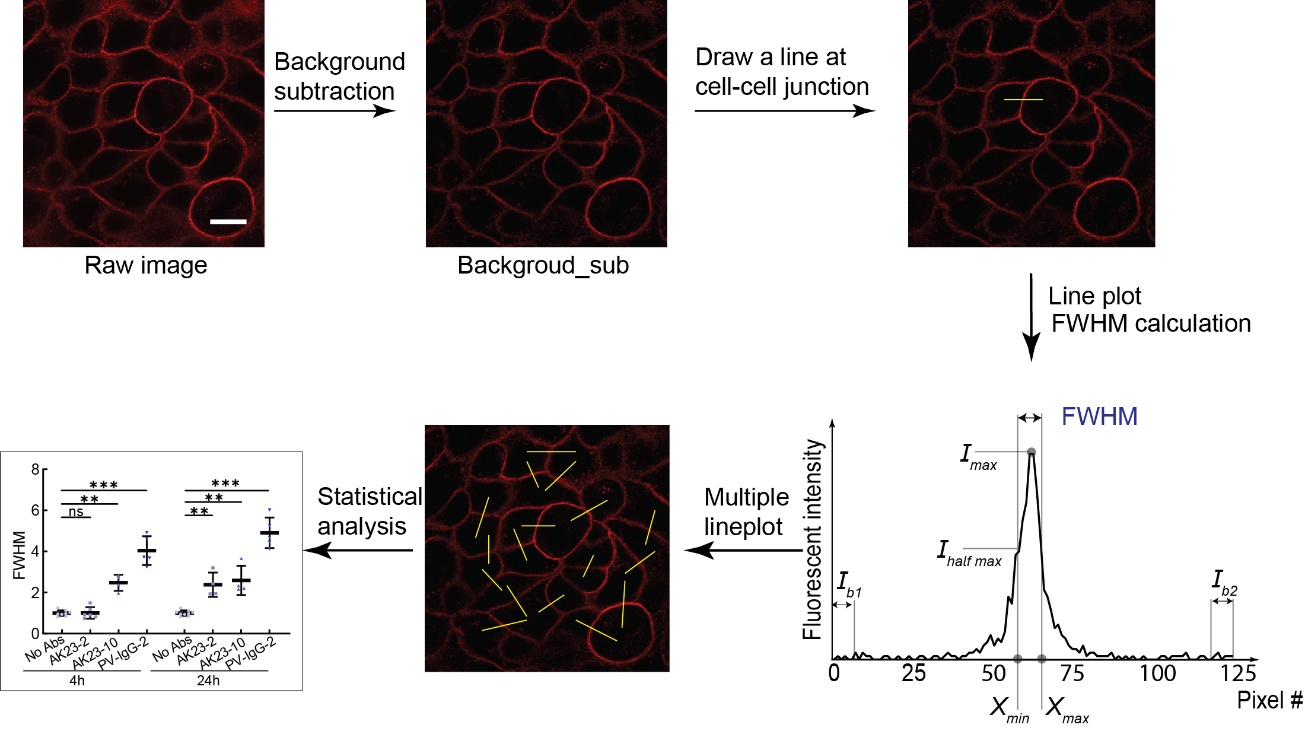

**Figure S1. Workflow of the image analysis for FWHM.**

The background was subtracted from the immunostaining images using the rolling ball in ImageJ. The brightness of background-subtracted images was adjusted in ImageJ to show representative images. In addition, a line was drawn at the cell-cell junction (from nucleus to nucleus) on the background-subtracted image. Plots were made with the fluorescence intensity *vs* pixel number along the reference line. I_max_: Absolute peak intensity; I_b_: Background noise that is defined as the average detected intensity of the first 5 pixels at each end of the reference line (I_b1_ and I_b2_); I_max_ - I_b_: Relative peak intensity; I_half max_: Half of the relative peak intensity; X_max_: Maximum pixel number where I_half max_ is reached; X_min_: Minimum pixel number where I_half max_ is reached; FWHM: X_max_ - X_min_. Statistical analysis was performed based on more than 10 images from three repeated samples and each image with at least 20 individual lines. Scale bar: 10 µm.

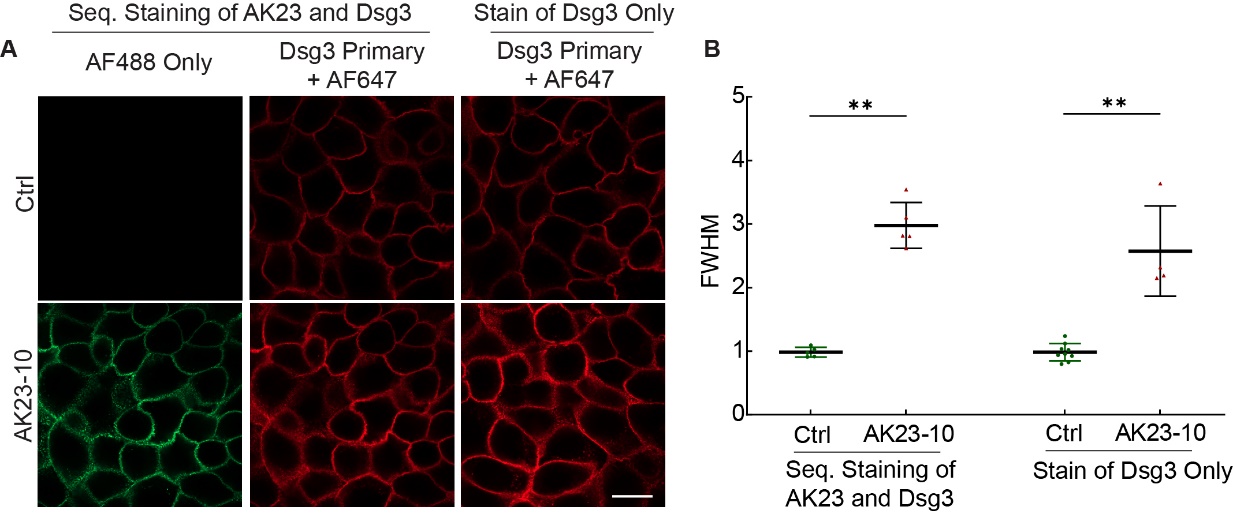

**Figure S2. Comparison of FWHM between sequential staining of AK23 and Dsg3 and staining Dsg3 only.**

HaCaT monolayers were stained in two ways: 1. Stained AK23 using anti-mouse Alexa Fluor 488 and then stained Dsg3 using anti-Dsg3 antibody followed by Alexa Flour 647; 2. Stained Dsg3 using anti-Dsg3 antibody followed by Alexa Flour 647. (A). Immunofluorescent images of two staining methods for both Control and AK23 (10 µg/mL, 24h). (B). FWHM analysis shows a negligible difference between the two staining methods. All values are mean ± SD (n≥10). * p<0.05, ** p<0.01, *** p<0.005, ns, p>0.05. Scale bar: 10 µm. Three repeats for all conditions.

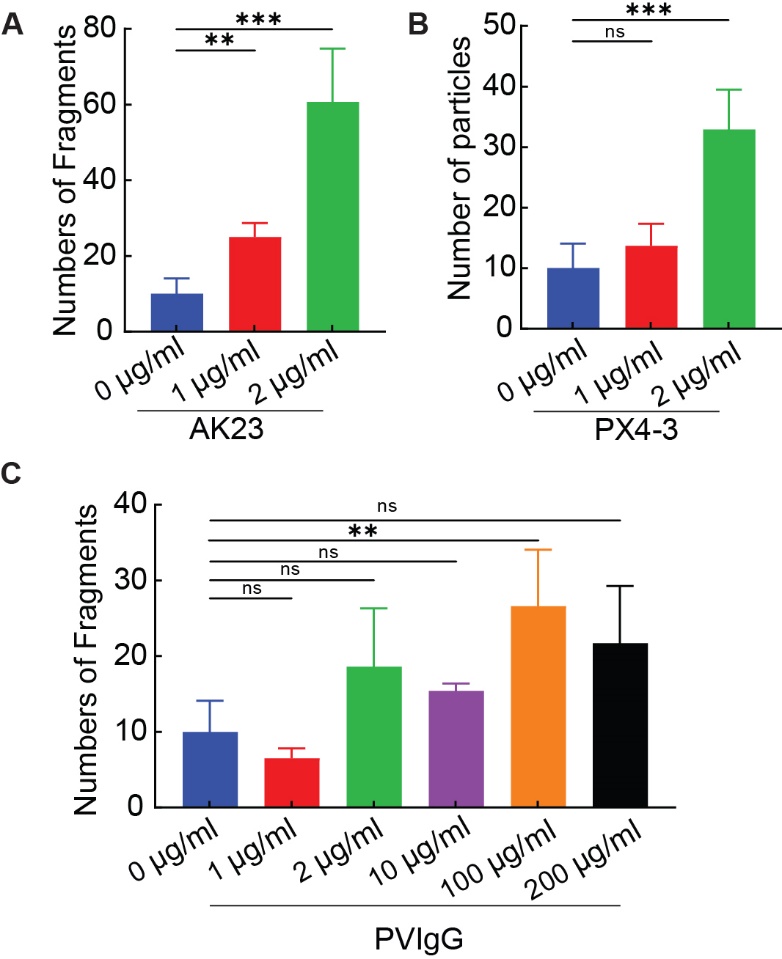

**Figure S3: Antibody titration experiments with keratinocyte dissociation assay (KDA).**

(A) The number of fragments for HaCaT monolayers in standard cell culture for two days treated with AK23 antibody at 1 μg/ml and 2 μg/ml compared to controls (no treatment). (B) The number of fragments for HaCaT monolayers treated with PX4-3 antibody at 1 μg/ml and 2 μg/ml compared to controls (no treatment). (C) The number of fragments for HaCaT monolayers treated with PX4-3 antibody at 1 μg/ml, 2 μg/ml, 10 μg/ml, 100 μg/ml, and 200 μg/ml as compared with controls (no treatment). All values are mean ± SD (n≥3). * p<0.05, ** p<0.01, *** p<0.005, ns, p>0.05. Scale bar: 10 µm. Three repeats for all conditions.

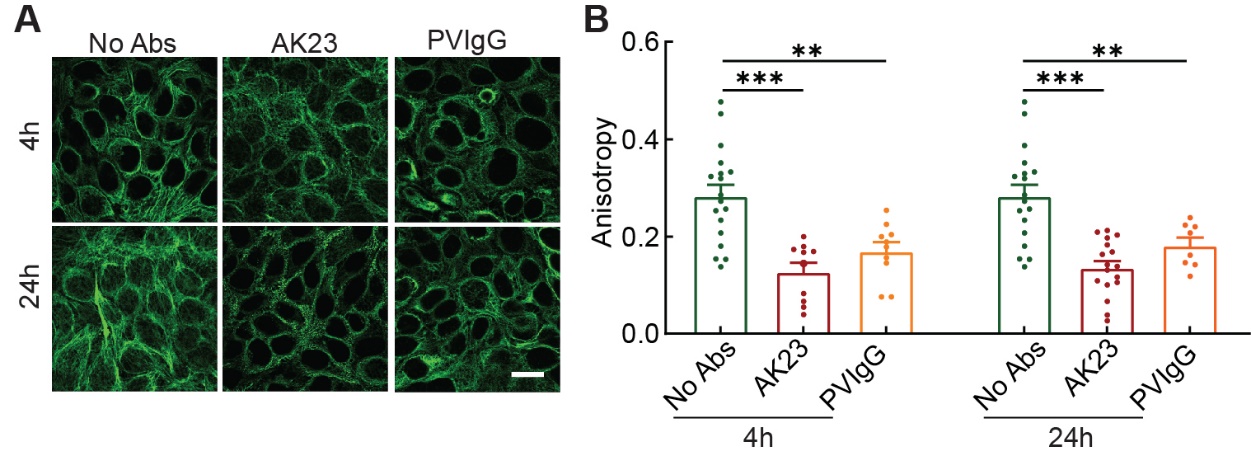

**Figure S4.** **PV Abs disordered the organization of intermediate filaments**

To evaluate the cytoskeletal activities induced by AK23 and PVIgG, we monitored the remodeling of the intermediate filament fibers via immunostaining. A customized MATLAB code was built for the anisotropy analysis. Briefly, background subtracted images were filtered by Butterworth filter that was built in MATLAB. The anisotropy was calculated by Fourier transform methods (FTM) ^1^. HaCaT cells were treated with 2 µg/mL of the following PV Abs: AK23 and PVIgG. Cells with no treatments were used as controls (No Abs). (A) Immunofluorescent images of intermediate filaments with AK23 and PVIgG treated at 4h and 24h. (B) Anisotropic analysis of intermediate filaments displays the retraction of intermediate filaments. All values are mean ± SD (n≥10). *p<0.05, **p<0.01, ***p<0.005, ns: p>0.05. Scale bar: 10 µm. Three repeats for all conditions.

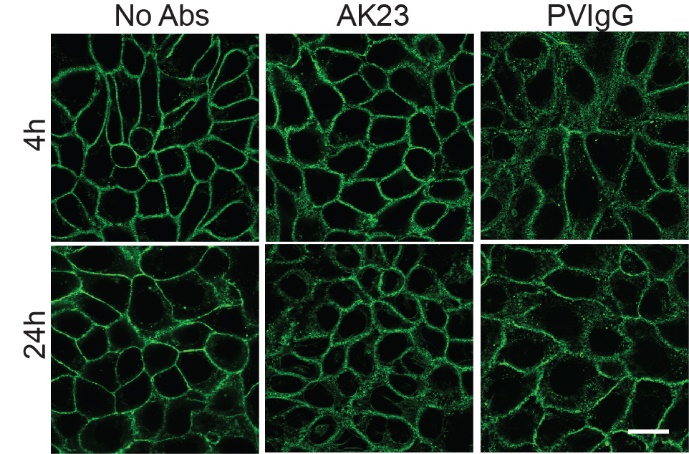

**Figure S5.** **PV Abs induced the remodeling of actin**.

HaCaT cells were treated with 2 µg/mL of the following PV Abs: AK23 and PVIgG. Cells with no treatments were used as controls (No Abs). Immunofluorescent images of actin with AK23 and PVIgG treated at 4h and 24h. Scale bar: 10 µm.

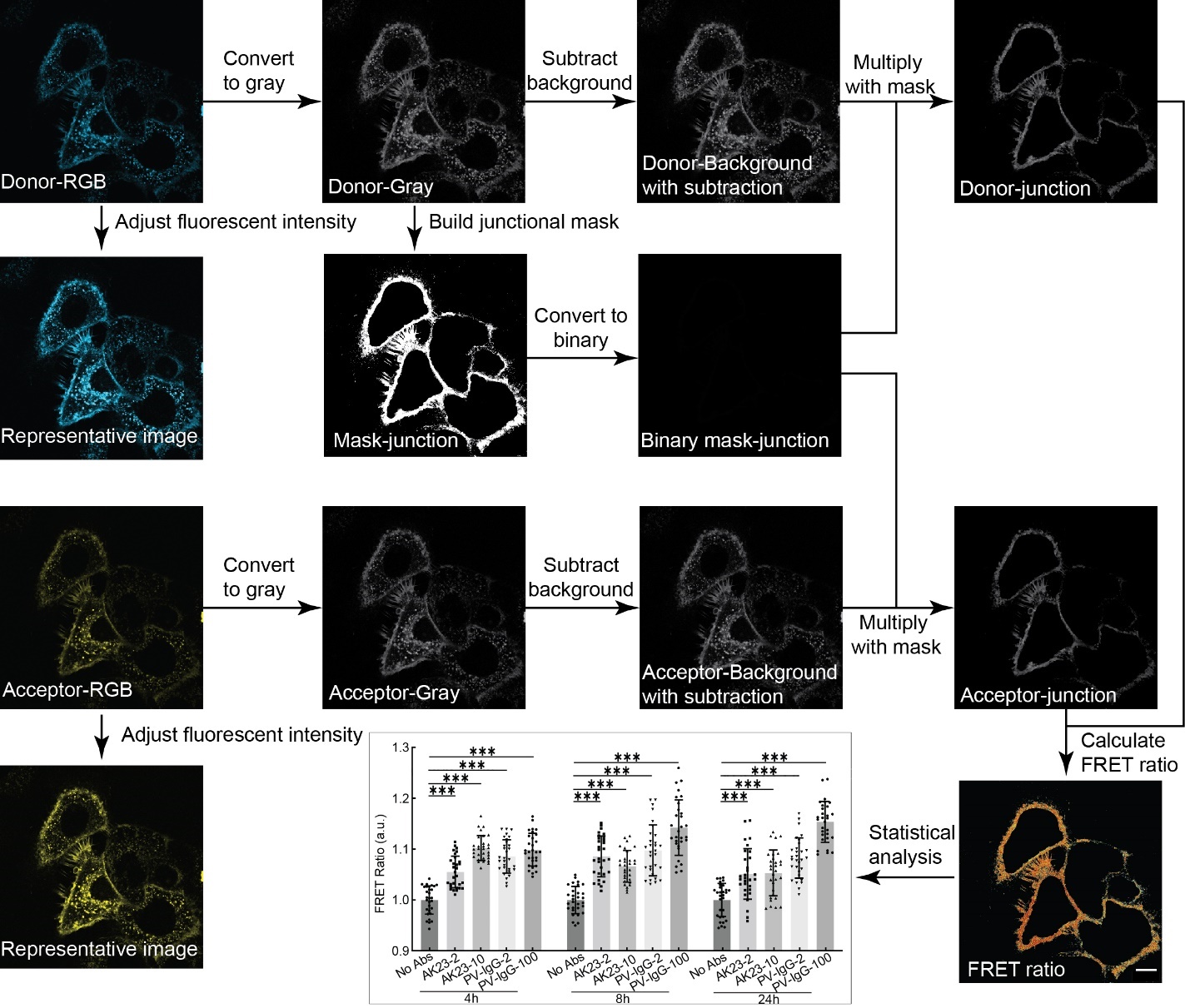

**Figure S6.** **Workflow of the Dsg3 FRET image process.**

To evaluate the tension changes and RhoA activation induced by PV Abs, FRET ratio was analyzed based on the fluorescent images. Figure S4 shows the workflow of the image process. Fluorescent images of the donor and acceptor were captured using the ZEISS confocal microscopy system. Before the quantitative image analysis, all RGB images were converted into gray-scale images, followed by background subtraction using the rolling ball in ImageJ. To focus on the FRET changes at cell-cell junctions, a mask outlining the junctions was built. Specifically, the noise in the cytoplasm was manually removed, followed by the adjustment of the threshold. The image was then converted into a binary mask. Background-subtracted images of the donor and acceptor were then multiplied with the mask. The FRET ratio was calculated by dividing the masked acceptor image by the corresponding masked donor image pixel by pixel. At least 30 image sets of donor and acceptor from three repeats were used for the FRET ratio analysis. A MATLAB script was built for the FRET ratio calculation and making the heatmap images. Statistical analysis and bar plot was performed in GraphPad Prism.­ Representative images were obtained by adjusting the brightness of RGB images. RGB images were converted into grayscale images and then subtracted from the background. A binary mask with cell junctions only was made from Donor-Gray by removing cytoplasmic noise and adjusting the threshold. The binary mask was then multiplied back to the donor with background subtraction and the acceptor with background subtraction to outline junctions. The FRET ratio was then calculated by dividing the Acceptor-junction image by the Donor-junction image pixel by pixel. Heatmap images of the FRET ratio were produced by plotting the FRET ratio pixel by pixel. At least 30 image sets from 3 repeated samples were processed for statistics. Statistical analysis and graph plot were performed via GraphPad Prism.

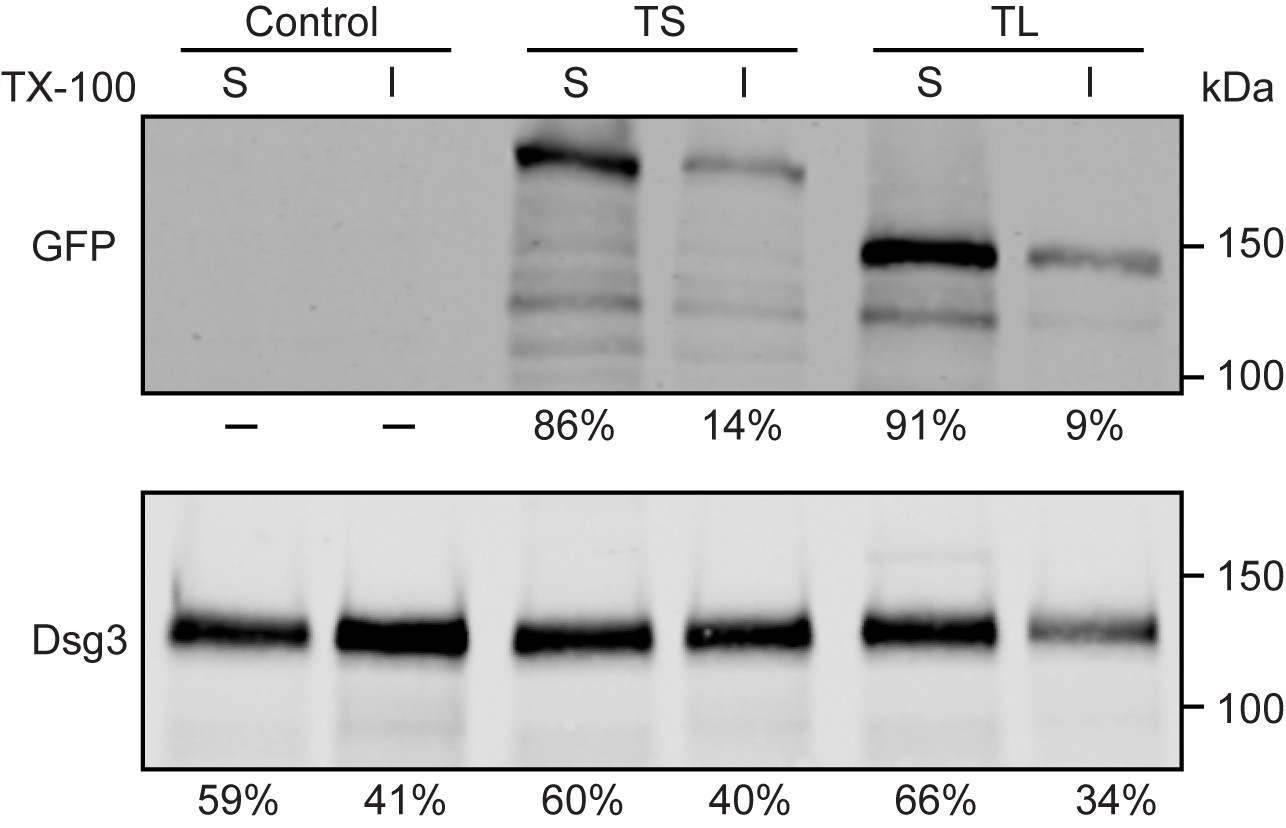

**Figure S7. Integration of the Dsg3 FRET sensor into desmosomes.**

To further investigate how the Dsg3 FRET sensors were integrated into the cell-cell contact, a Triton X-100 fractionation experiment was performed to find the fraction of the sensors that were integrated into fully formed desmosomes and unbounded on the plasma membrane. GFP was stained to quantify the expression of the Dsg3 FRET sensors, and Dsg3 was stained to quantify the expression of endogenous Dsg3. The percentage of the total protein in the soluble (S) and insoluble (I) fractions is shown below the blots.

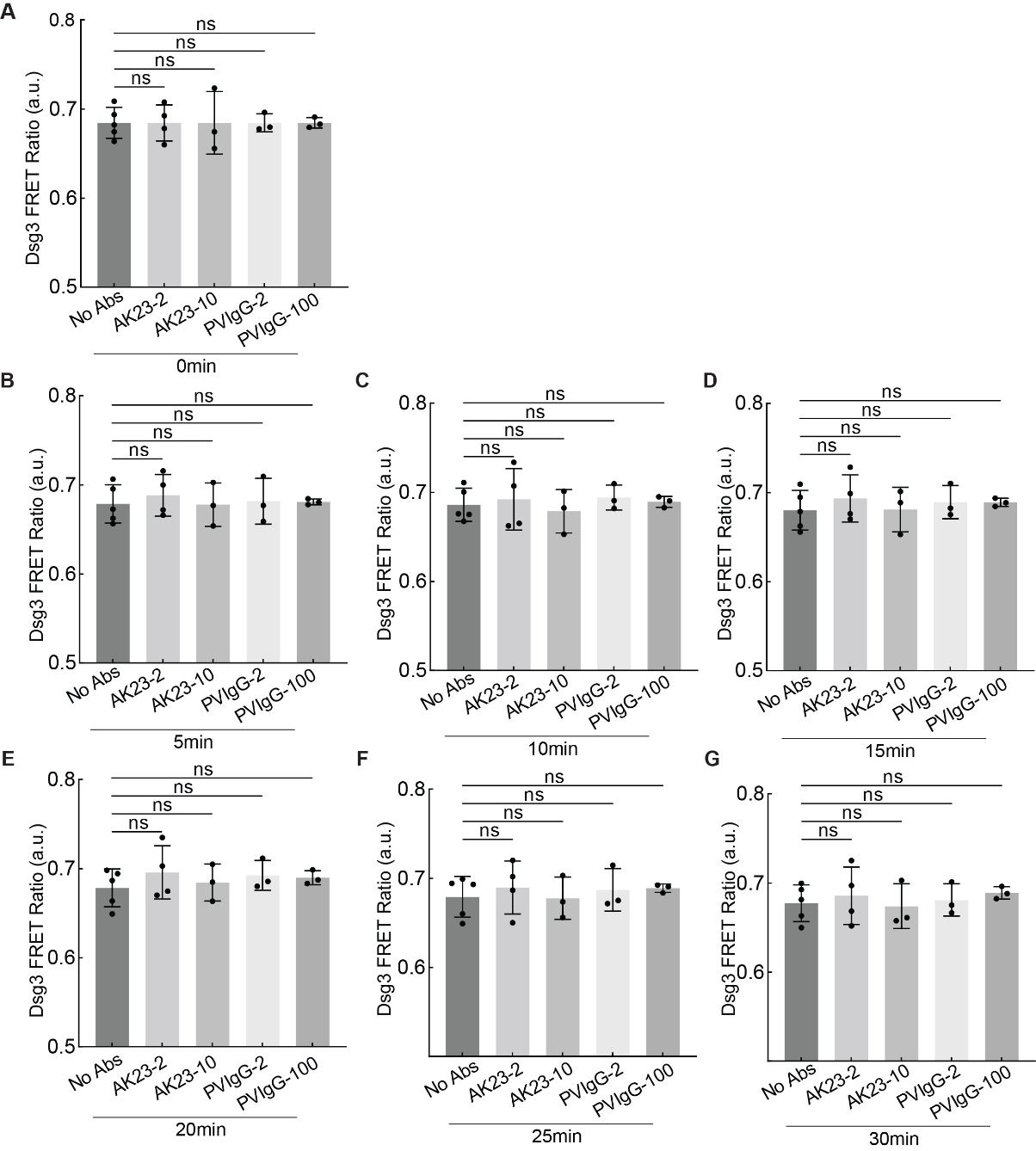

**Figure S8. Short-term observation of Dsg3 FRET ratio.**

To evaluate how PV Abs induce tension changes in Dsg3 in the short-term, we monitored the Dsg3 FRET ratio in real time. For 30 minutes HaCaT cells were treated with AK23 at 2 μg/mL (AK23-2), AK23 at 10 μg/mL (AK23-10), PVIgG at 2 μg/mL (PVIgG-2), and PVIgG at 100 μg/mL (PVIgG-100). The controls were performed with no antibody treatment (No Abs). The FRET ratio was calculated by dividing the acceptor by the donor. Tracking Dsg3 FRET over the first 30 min of treatment with these antibodies reveals no significant change in the short-term. One image was taken every 5 minutes. All values are mean ± SD. Three repeats for all conditions.

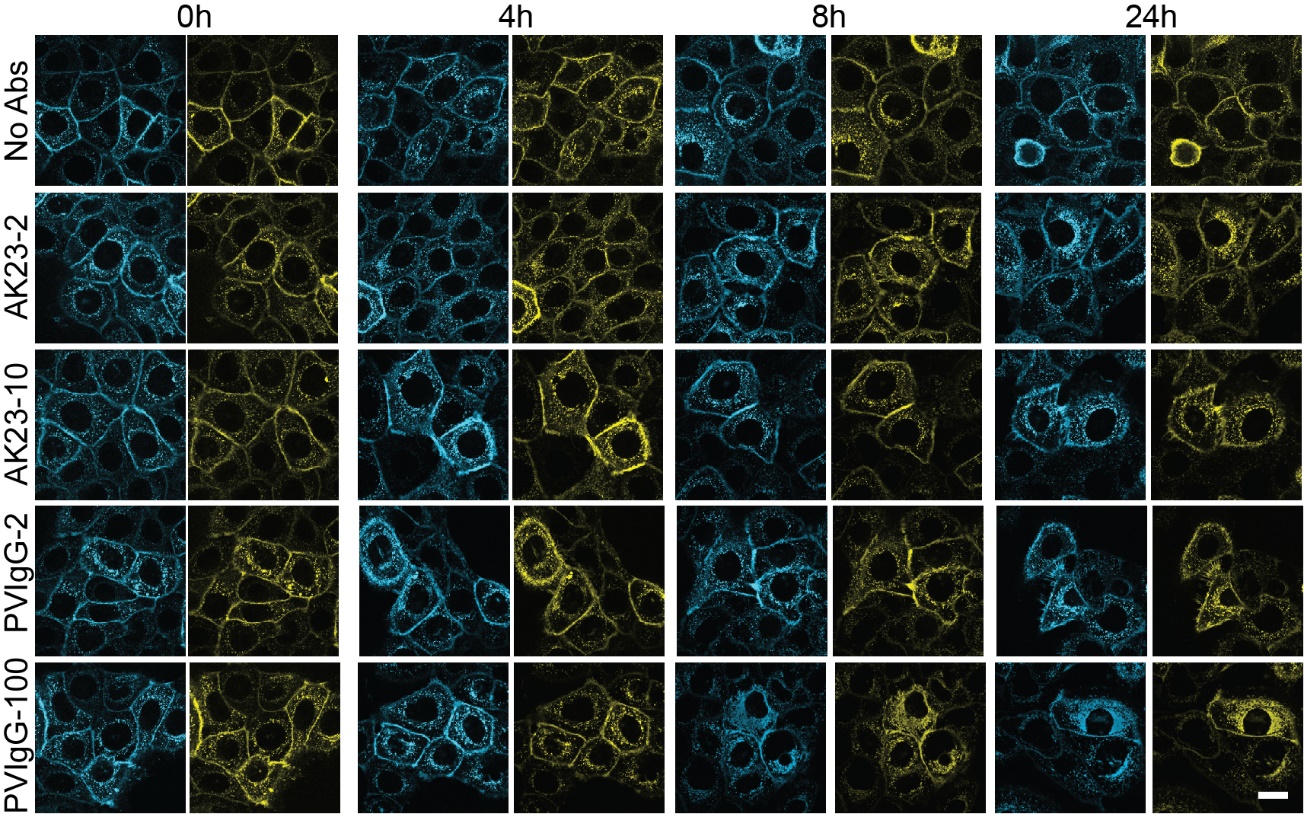

**Figure S9. Fluorescent images of Dsg3 FRET sensor**.

Here, HaCaT monolayers were exposed to a commercial mouse-derived monoclonal antibody (AK23) and patient-derived polyclonal antibody (PVIgG) produced by purifying the serum from patients with PV diseases. For AK23, 2 μg/mL and 10 μg/mL were added to HaCaT monolayers. In addition, PVIgG with 2 μg/mL and 100 μg/mL was used to study the different effects between the two types of antibodies. Fluorescent images of all antibody-treated groups were captured at pre-treated (0h) and post-treated (4h, 8h, and 24h) time points. Fluorescent images of both donor (mTFP, blue) and acceptor (Venus, yellow) channels were obtained in HaCaT cells at the time points of 0h, 4h, 8h, and 24h. Cells were treated in four conditions: no antibody treatment (No Abs), AK23 at 2 μg/mL (AK23-2), AK23 at 10 μg/mL (AK23-10), PVIgG at 2 μg/mL (PVIgG-2) and PVIgG at 100 μg/mL (PVIgG-100). Scale bar: 10 µm.

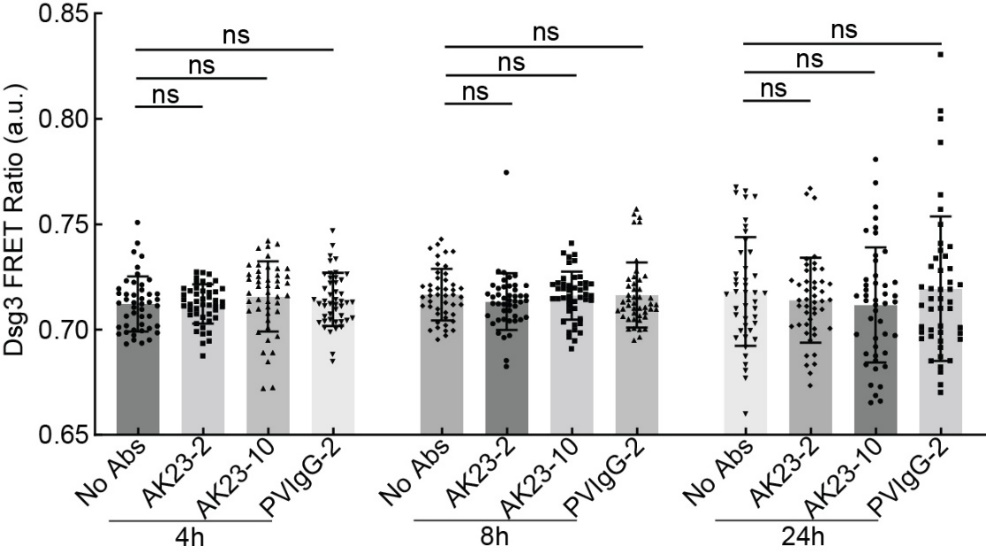

**Figure S10. PV Abs induce no FRET ratio changes on Dsg3 tailless control sensor.**

HaCaT cells were treated with AK23 at 2 μg/mL (AK23-2), AK23 at 10 μg/mL (AK23-10), and PVIgG at 2 μg/mL (PVIgG-2). The controls were performed with no antibody treatment (No Abs). The FRET ratio was calculated by dividing the acceptor by the donor. All values are mean ± SD. Three repeats for all conditions.

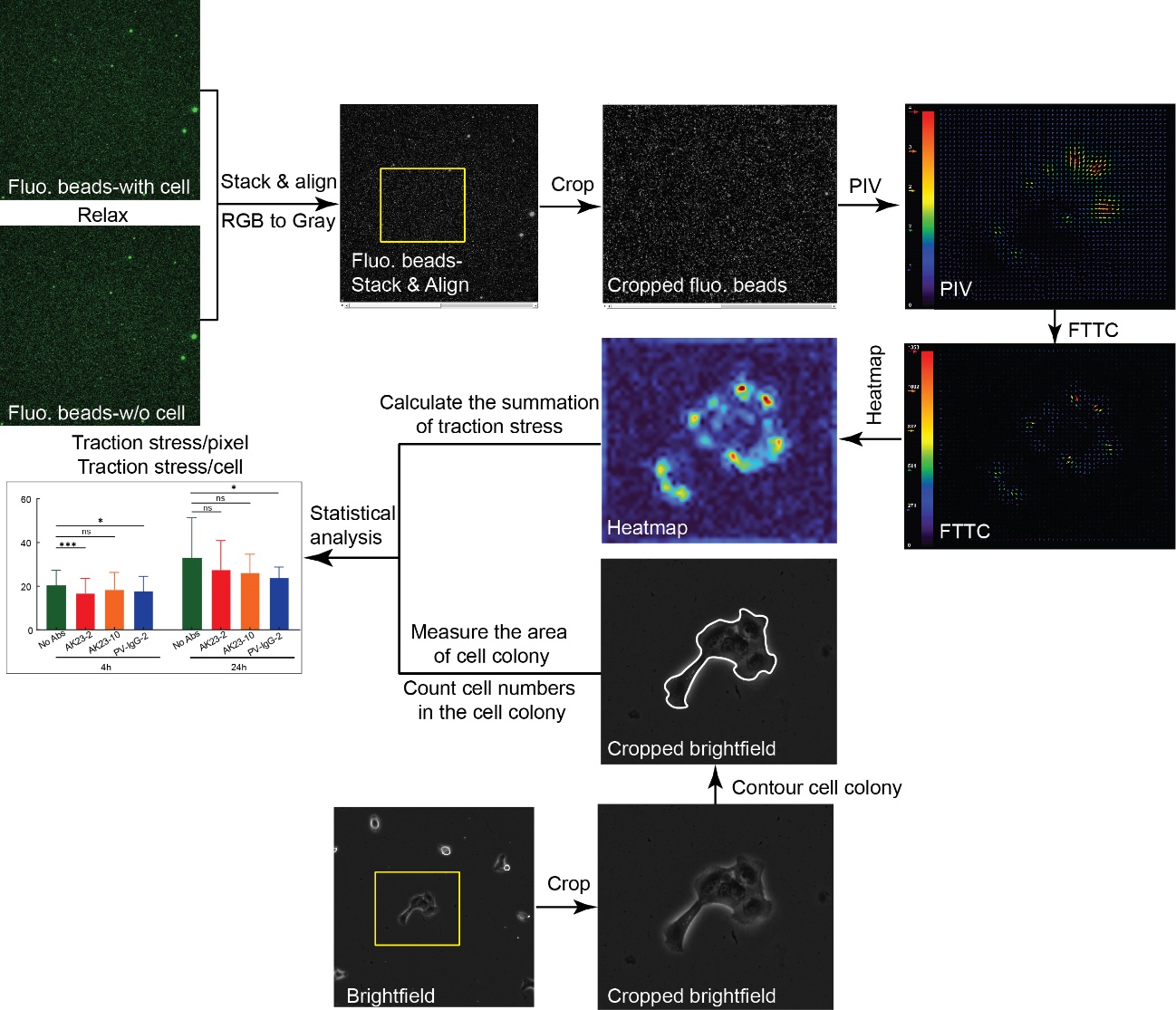

**Figure S11.** **Workflow for** **processing the TFM images.**

Traction force microscopy (TFM) was used to evaluate the traction stress and force changes at the cell-ECM junction. Fluorescent beads images with cell attachment (tense) and without cell attachment (relax) were collected using Lecia microscopy system, stacked, and aligned in ImageJ. The ROI of the cell colony was selected based on the brightfield image and then was applied to the beads image to crop the corresponding stacked fluorescent beads (Fluo. beads) image. The displacement of beads was then analyzed using the PIV plugin, followed by traction stress analysis via the FTTC plugin in ImageJ. A customized MATLAB script was used to calculate traction force and plot heatmap images. The area of the cell colony was calculated by manually drawing the outline of the cell colony and measured in ImageJ. Cell numbers were also counted manually. Traction stress/pixel was calculated by dividing the summation of traction stress by the area. Traction stress/cell was calculated by dividing the summation of traction stress by the cell numbers of the colony. At least 10 images from three repeats were used for the statistical analysis.

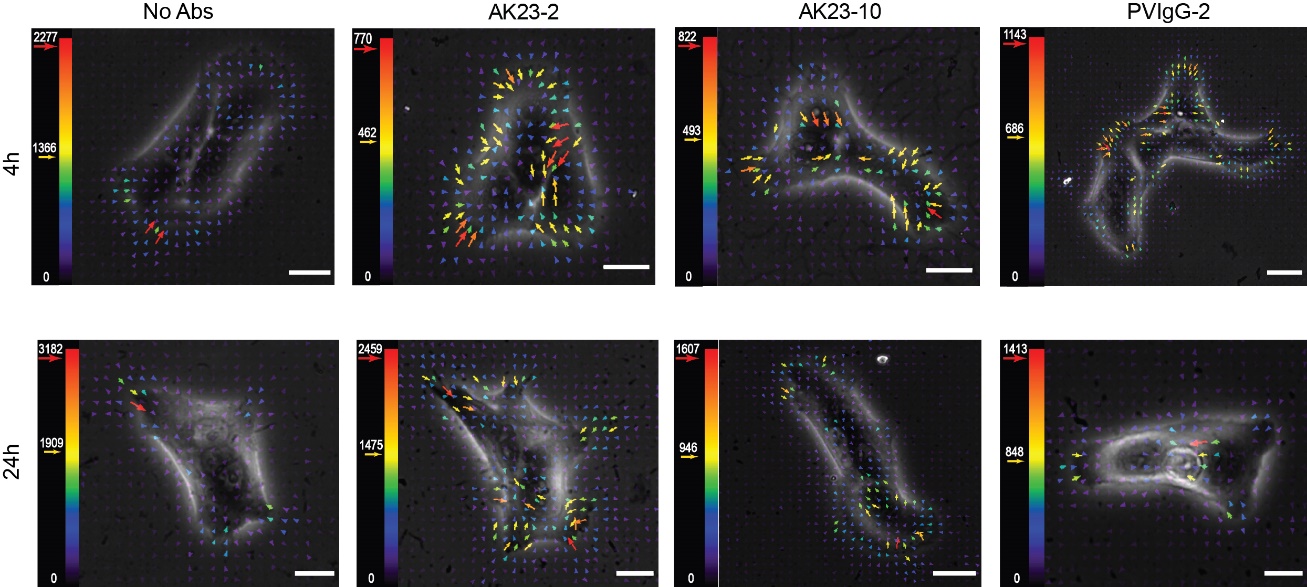

**Figure S12.** **Overlapped stress and brightfield.**

HaCaT cells were seeded on gels coated with 0.2 μm fluorescent beads overnight to let them attach to gels. Tense and relaxed fluorescent beads images were taken and analyzed through the workflow shown in Figure S8. Cells were treated with four different conditions: No antibody treatment (No Abs), 2 μg/mL of AK23 (AK23-2), 10 μg/mL of AK23 (AK23-10) and 2 μg/mL of PVIgG (PVIgG-2). Fluorescent image sets of beads were taken at both 4h and 24h for each experimental group. The color of the arrow represents the magnitude of the traction stress vector. Scale bar: 15 µm.

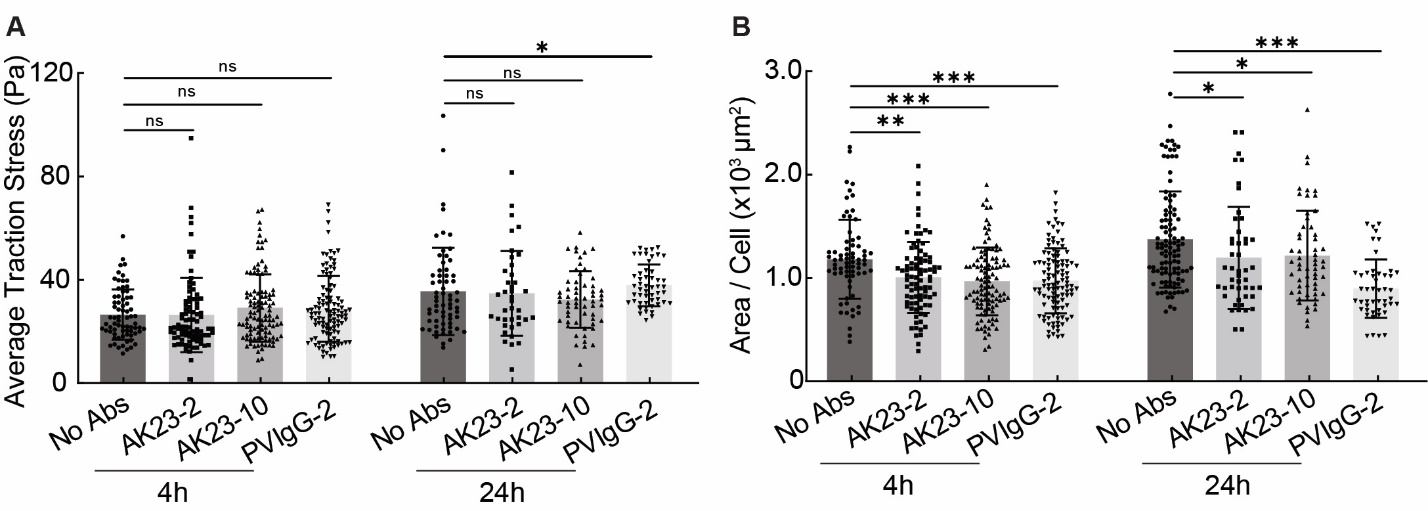

**Figure S13.** **Average traction stress and area per cell in TFM experiments.**

The average traction stress was calculated by dividing the summation of traction stress of the image by the area of the cell colony, and the area per cell was calculated by dividing the area of the cell colony by the number of cells in the colony. The area of the cell colony was manually outlined and measured in ImageJ. (A) Average traction stress at 4h and 24h. (B) Area per cell at 4h and 24h. All values are mean ± SD (n≥10). * p<0.05, **p<0.01, ***p<0.005, ns: p>0.05. Three repeats for all conditions.

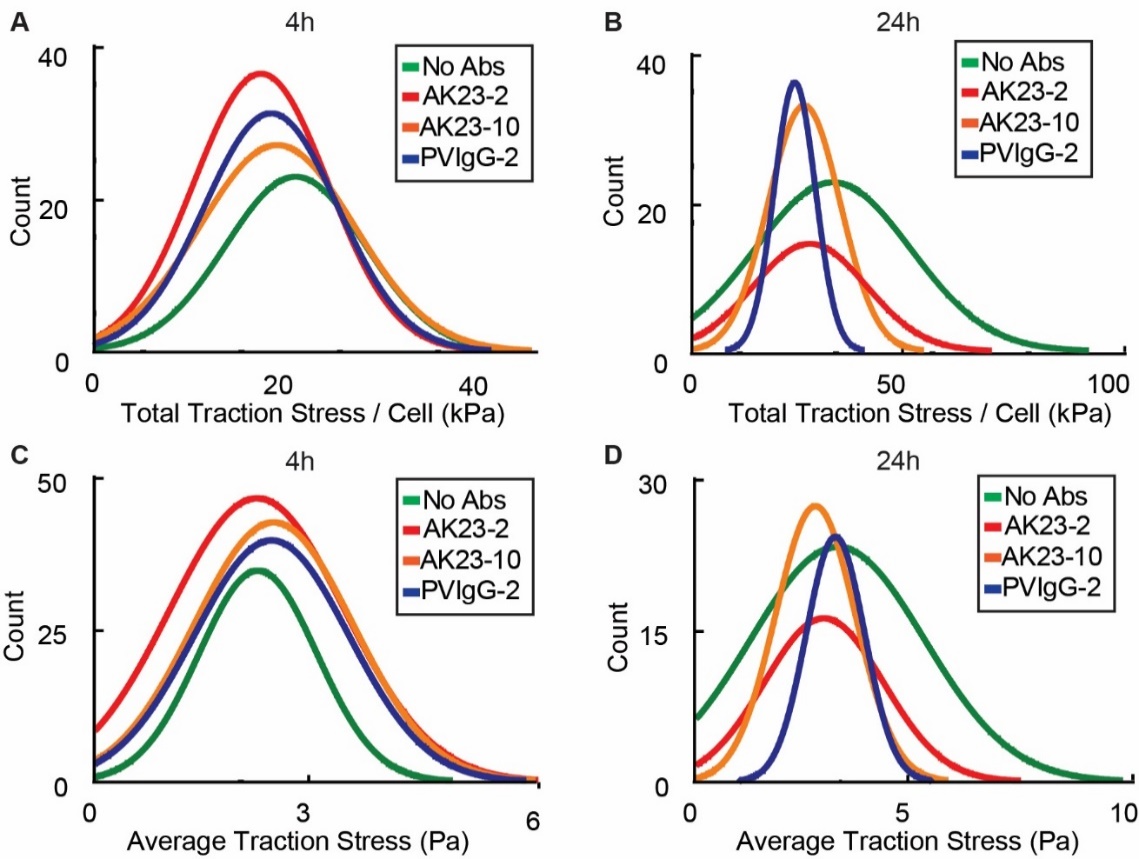

**Figure S14. Distribution of traction stress/pixel and traction stress/cells**.

(A) Traction stress/cell at 4h. (B) Traction stress/cell at 24h. (C) Traction stress/pixel at 4h. (D) Traction stress/pixel at 24h

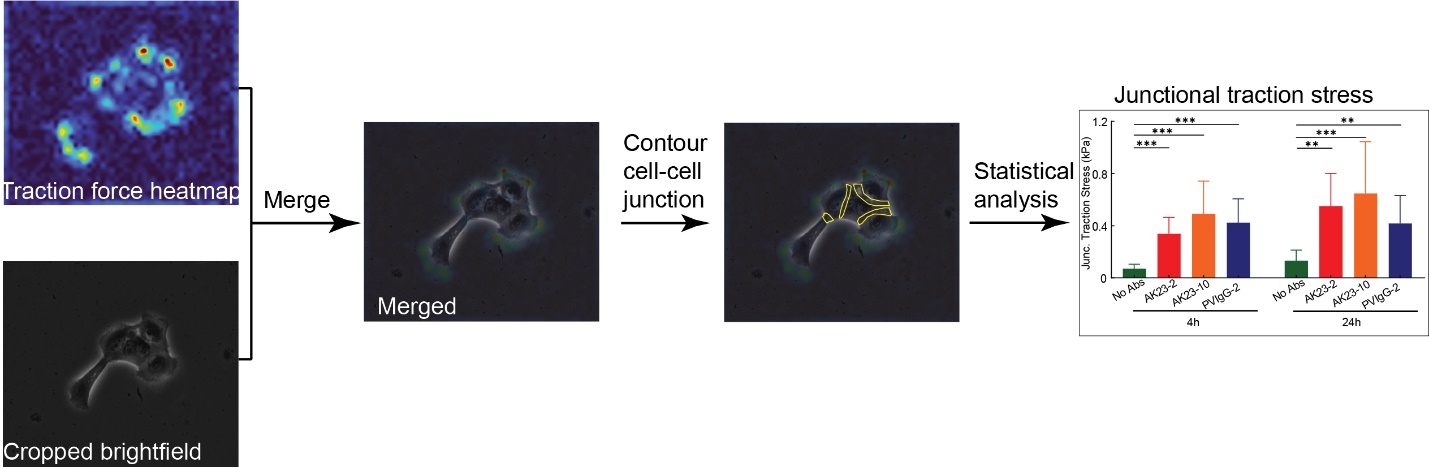

**Figure S15.** **Workflow of junctional traction stress calculation.**

To calculate the junctional traction stress, the heatmap image of traction stress and cropped brightfield image were obtained using the method shown in Figure S8. Two images overlapped to assist with distinguishing the cell-cell junctions. A customized MATLAB script was used to contour the cell-cell junctions. At least 10 images from three repeats were used for statistical analysis. Cell-cell junction was manually outlined by picking at least 10 location points around the junction which would form a closed region on the brightfield image. The intensity at each pixel of the traction stress images in the same region was then extracted and averaged for statistical analysis.

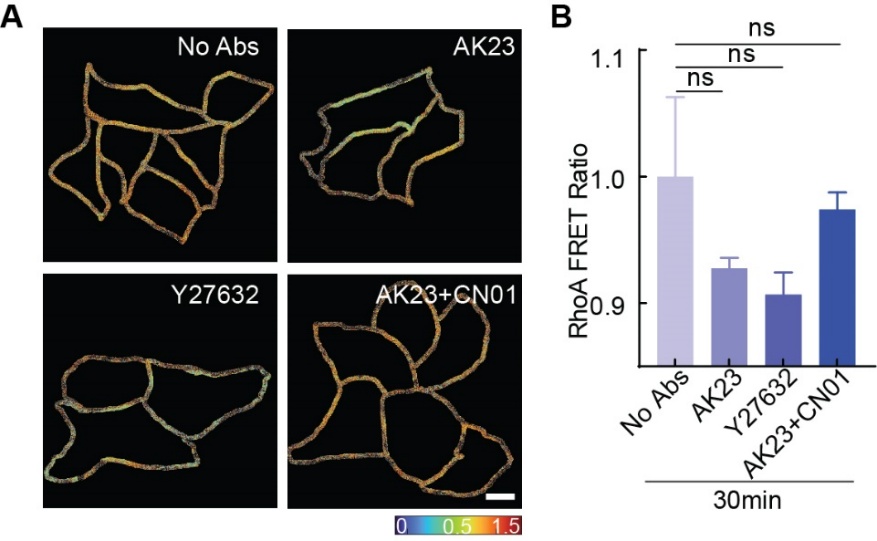

**Figure S16.** **RhoA FRET ratio in response to different treatments**.

HaCaT cells were treated with 2 µg/mL AK23, 10 µM ROCK inhibitor (Y27632), and 2 µg/mL AK23 with 1 unit/mL CN01. (A) Heatmap images of RhoA FRET sensor with different treatments. (B) Quantitative analysis of the RhoA FRET ratio. At least 6 image sets from three repeats were used for statistical analysis. All values are mean ± SD. *p<0.05, **p<0.01, ***p<0.005, ns: p>0.05. Three repeats for all conditions. Scale bar: 10 µm.

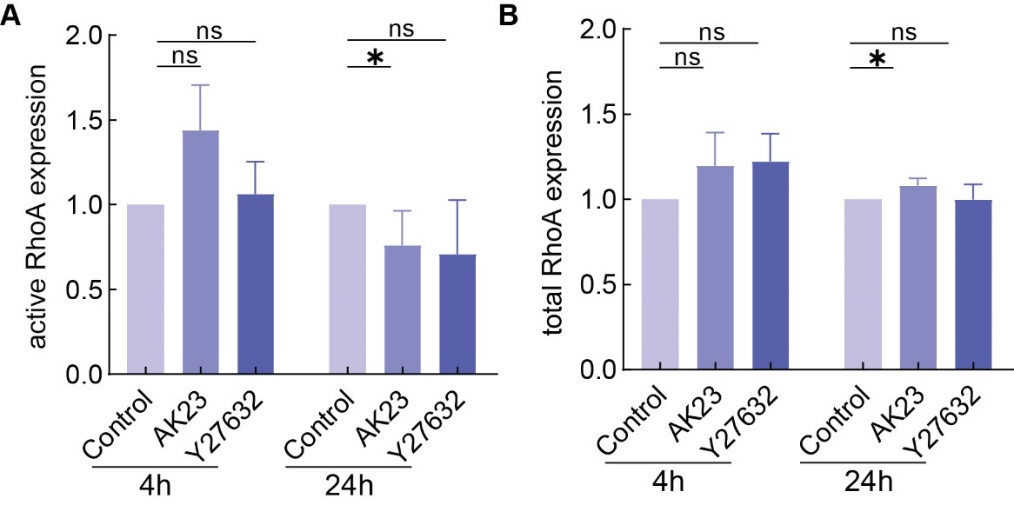

**Figure S17.** **Western blot results display the activities of RhoA regulated by AK23**.

(A) Quantitative results of western blot for active RhoA expression induced by 2 µg/mL of AK23 and 10 µM of Y27632. (B) Quantitative results of western blot for total RhoA expression induced by 2 µg/mL of AK23 and 10 µM of Y27632.

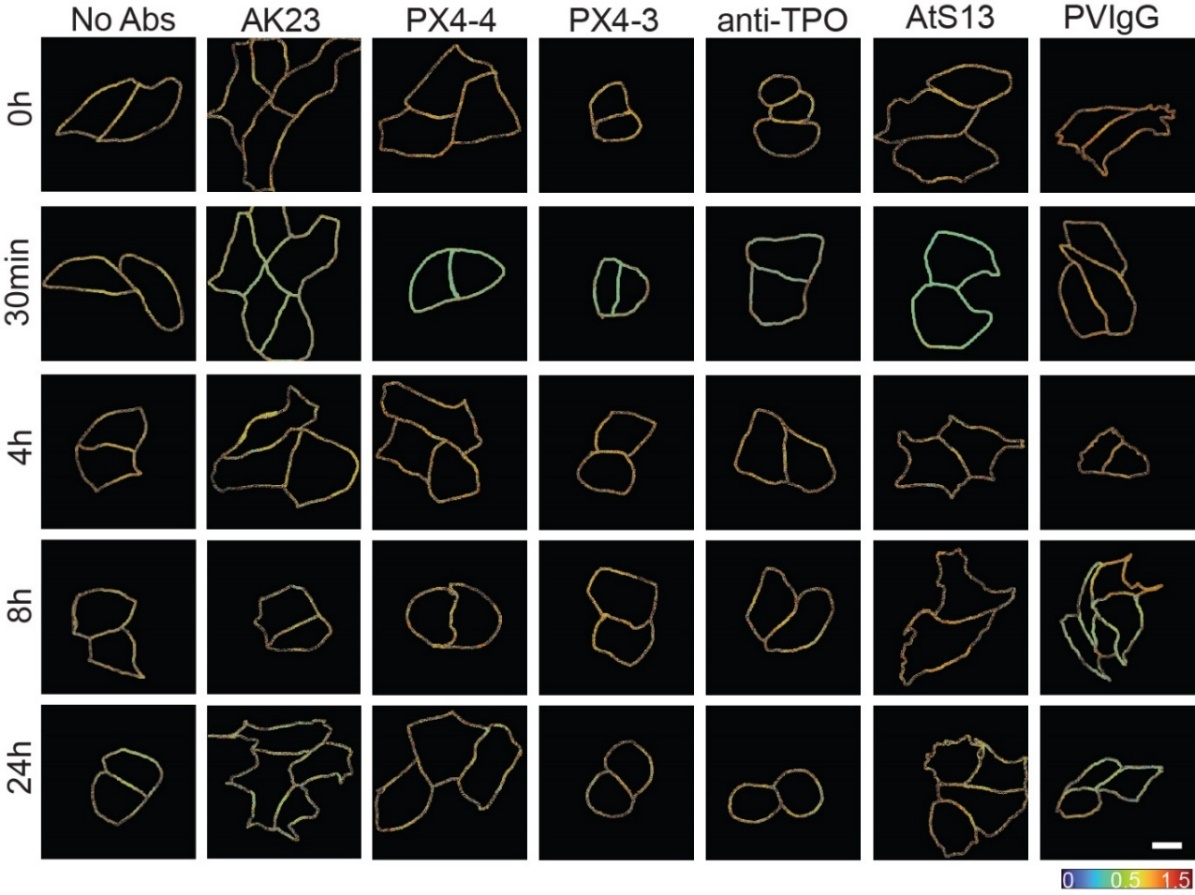

**Figure S18.** **Decreasing RhoA FRET ratio induced by PV Abs**.

To investigate the effects of PV Abs on the activities of active RhoA, we observed the activation of RhoA using the RhoA FRET sensor. Similar to the Dsg3 FRET sensor, only junctional RhoA was evaluated. Different antibodies, including AK23, PX4-4, PX4-3, anti-TPO, AtS13, and PVIgG, were added into HaCaT monolayers with a concentration of 2 µg/mL individually. Using the image process mentioned in Figure S4, heatmap images were calculated. Heatmap images of RhoA FRET ratio at cell-cell junction displayed the reduced activation of RhoA at cell-cell contacts treated with 2 µg/mL of AK23, PX4-4, PX4-3, anti-TPO, AtS13, and PVIgG. Cells with no antibody treatments served as control (No Abs). Scale bar: 10 µm.

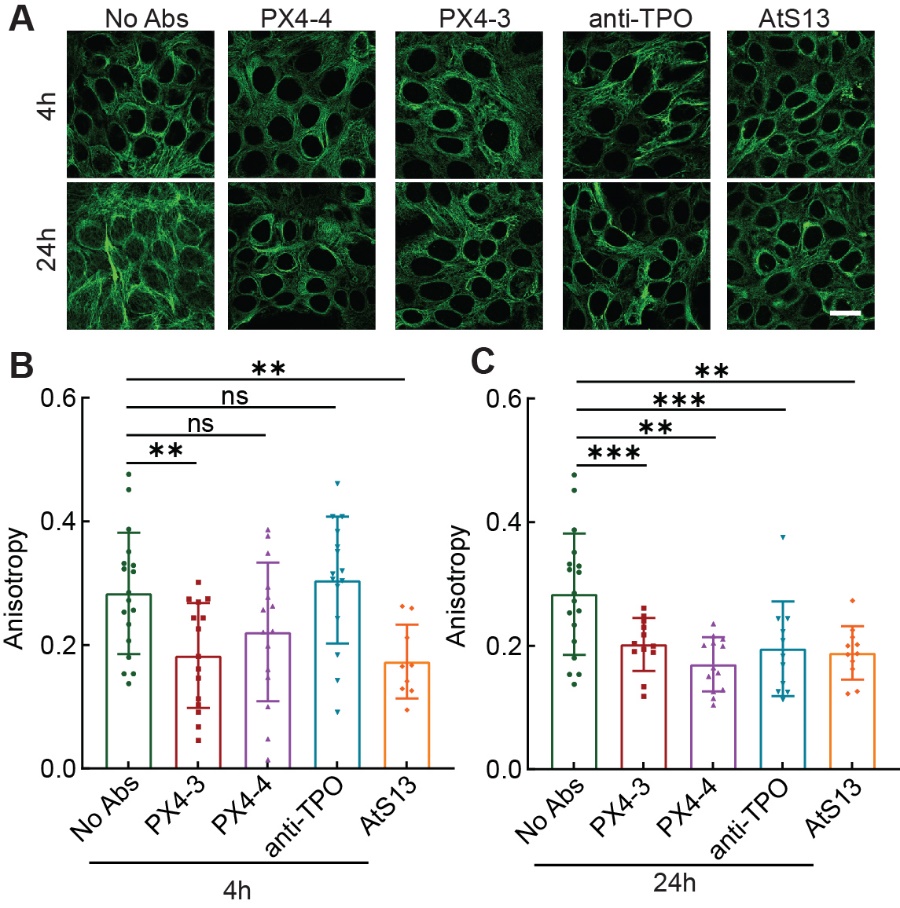

**Figure S19.** **Retraction of intermediate filaments induced by PV Abs**.

HaCaT cells were treated with 2 µg/mL of the following PV Abs: PX4-4, PX4-3, anti-TPO, and AtS13. Cells with no treatments were served as blank controls (No Abs). (A) Immunofluorescent images of intermediate filaments with PV Abs treated at 4h and 24h. (B, C) Anisotropic analysis of intermediate filaments displays the retraction of intermediate filaments. All values are mean ± SD (n≥10). *p<0.05, **p<0.01, ***p<0.005, ns: p>0.05. Scale bar: 10 µm. Three repeats for all conditions.

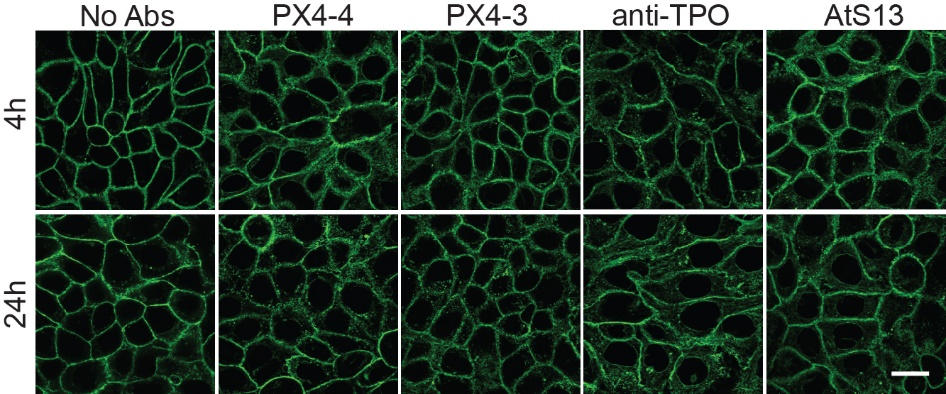

**Figure S20.** **PV Abs induced the remodeling of actin**.

HaCaT cells were treated with 2 µg/mL of the following PV Abs: PX4-4, PX4-3, anti-TPO, and AtS13. Cells with no treatments were served as blank controls (No Abs). Immunofluorescent images of actin with PX4-4, PX4-3, anti-TPO, and AtS13 treated at 4h and 24h. Scale bar: 10 µm.

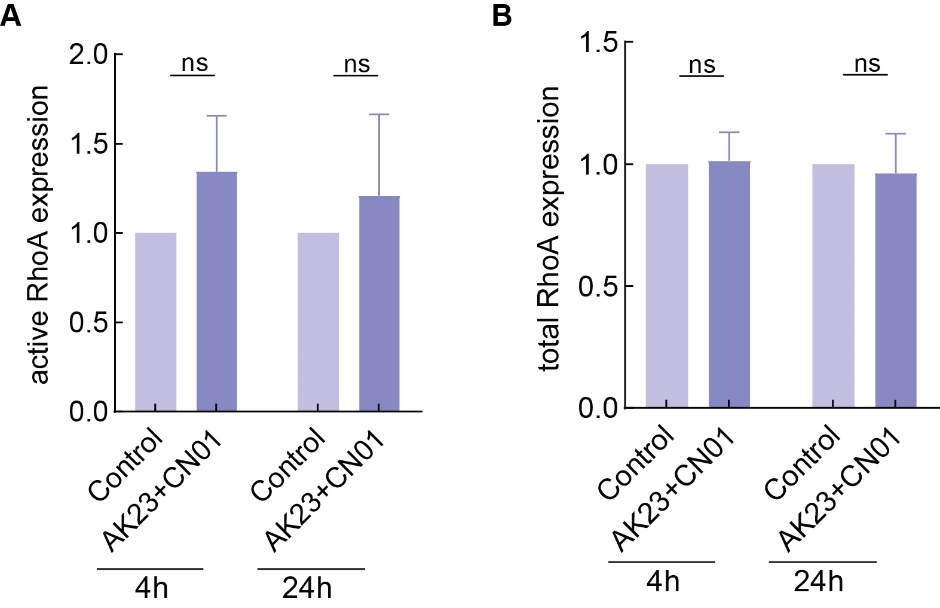

**Figure S21. Western blot results display the activities of RhoA regulated by CN01**.

(A) Quantitative results of western blot for active RhoA expression induced by 2 µg/mL of AK23 with 1 unit/mL CN01. (B) Quantitative results of western blot for total RhoA expression induced by 2 µg/mL of AK23 with 1 unit/mL CN01.

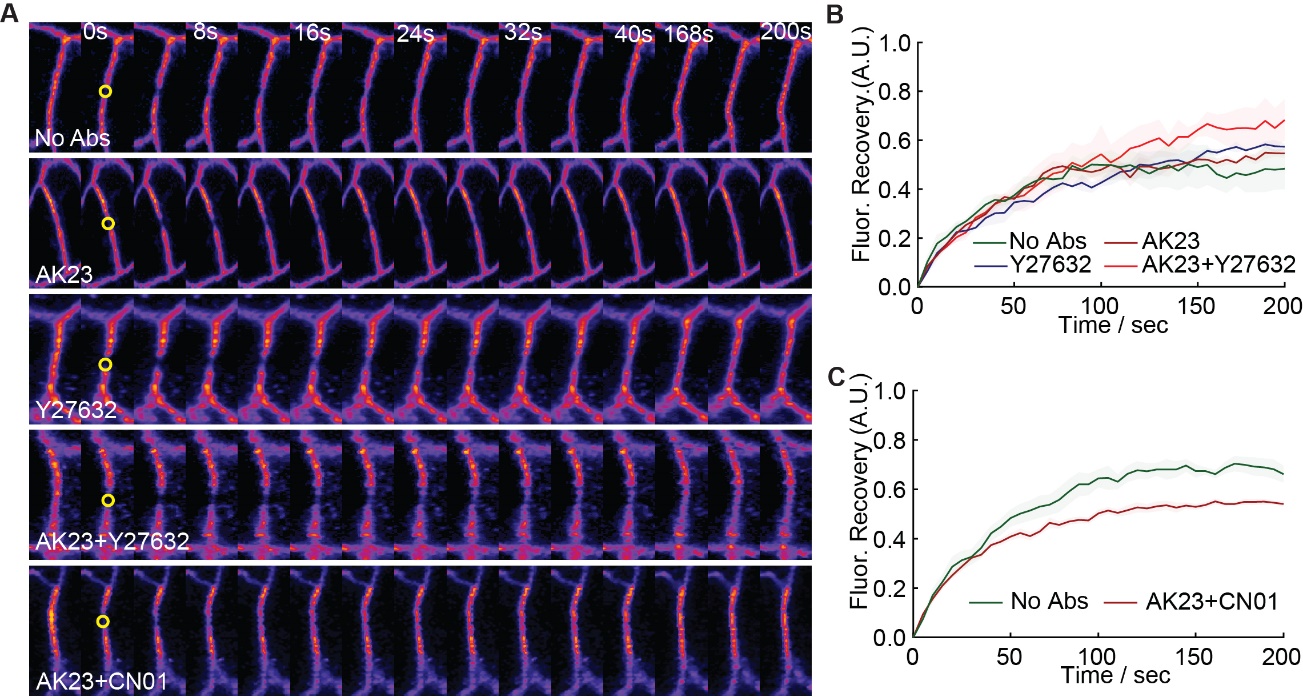

**Figure S22. FRAP of E-cadherin with different treatments**.

HaCaT cells were engineered to express E-cad–GFP. The dynamics of E-cad were observed using Fluorescence Recovery after Photobleaching (FRAP). Specifically, HaCaT cells were cultured to 80% confluency and were imaged using a ZEISS confocal microscopy system equipped with photobleaching and time-lapse modules. To obtain the recovery curve, an ROI with a diameter of 2 µm was drawn on the E-cadherin junction for photobleaching. In addition, an ROI with the same dimension was drawn on the black area as the background. Further, a reference signal was obtained with an ROI on a region of the E-cad junction without photobleaching. At least 5 images were captured before the photobleaching to measure the starting level. Post-bleached images were obtained every 8 seconds until no obvious fluorescent changes in the bleached area were observed. Here, the end time point was 200s. Collected data were calibrated and normalized using the easyFRAP-web ^2^. Cells were treated with 2 µg/mL AK23, 10 µM Y27632, 2 µg/mL AK23 plus 10 µM Y27632, and 2 µg/mL AK23 coupled with 1 unit/mL CN01 for 4 hours. (A) Montage images of the bleached E-cad junction were built by cropping the bleached junction from the observed fluorescent images at multiple selected time points and organizing them into montage templates in ImageJ. (B) FRAP recovery curve for cells treated with AK23 and Y7632. (C) FRAP recovery images for cells treated with AK23 and CN01. Three repeats for all conditions.

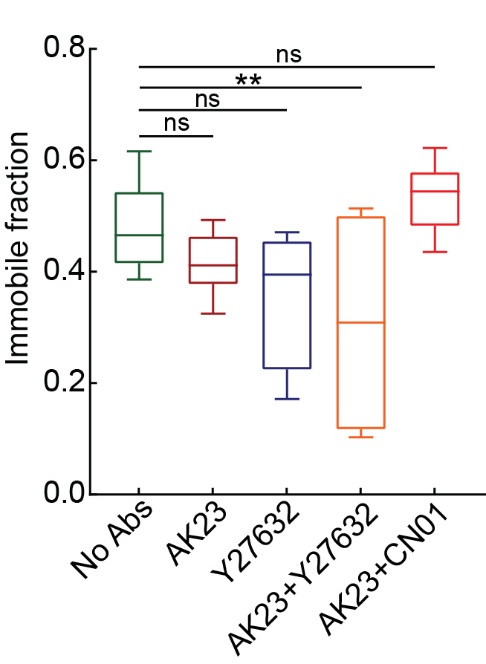

**Figure S23.** **Immobile fraction analysis.**

The corrected FRAP recovery curve was fitted by the nonlinear fitting model built in GraphPad Prism 9. A mobile fraction was directly extracted from the fitting result. The immobile fraction was calculated by subtracting the mobile fraction from 1. At least 5 separate FRAP data were captured for each sample, and 3 repeats were performed for each treated group. All values are mean ± SD. *p<0.05, **p<0.01, ***p<0.005, ns: p>0.05. Three repeats for all conditions.

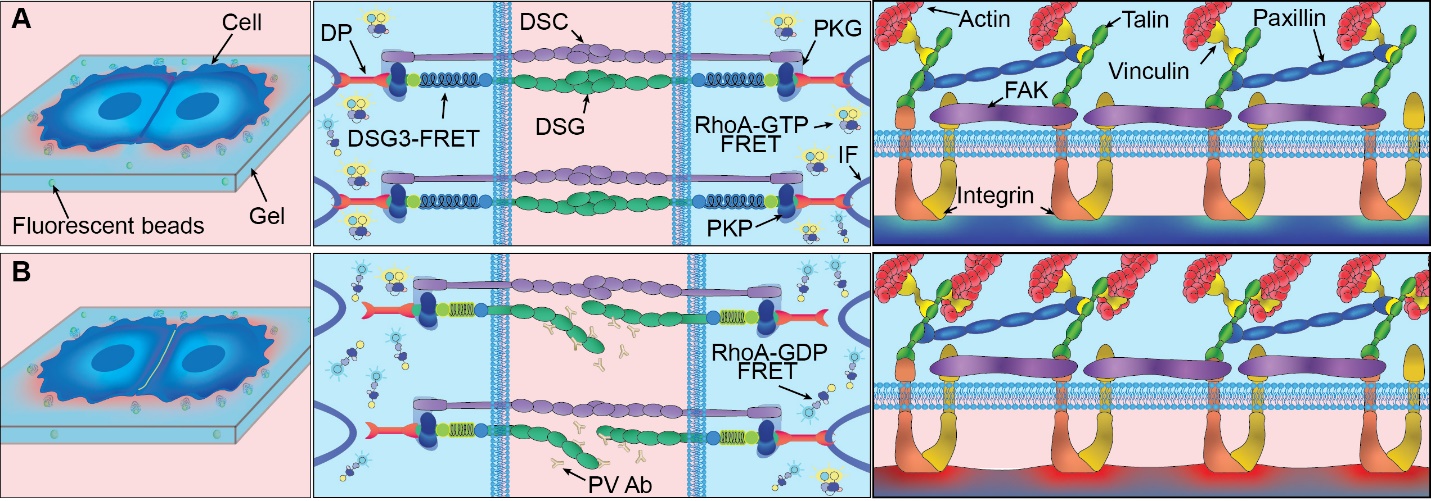

**Figure S24. Diagram of changes to keratinocytes under influence of PV-associated antibodies.**

(A) Under normal conditions, cells exert traction stress at the periphery of colonies, desmosomes and AJs remain intact, and focal adhesions underneath the cell-cell junction maintain lower forces. (B) Under exposure to PV Abs, traction stress beneath the cell-cell junction is elevated, linkages within desmosomes and AJs are disrupted, and focal adhesions underneath the cell-cell junction experience elevated tension.

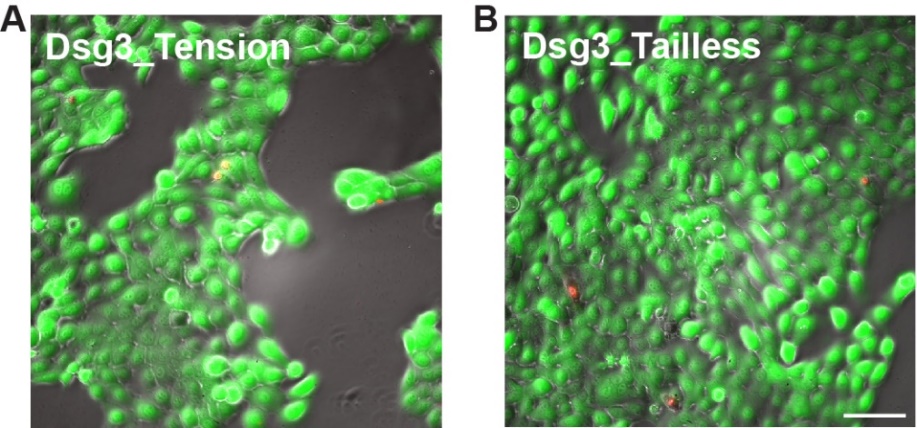

**Figure S25. Viability of HaCaT cells expressed Dsg3 FRET sensor.**

(A) Dsg3 tension sensor expressed HaCaT cells. (B) Dsg3 tailless sensor expressed HaCaT cells.
